# Biophysical mechanisms of default mode network function and dysfunction

**DOI:** 10.1101/2025.04.16.649208

**Authors:** Trang-Anh E. Nghiem, Vinod Menon

## Abstract

The default mode network (DMN) plays a fundamental role in internally focused cognition, and its disruption is implicated in numerous brain disorders. While neuroimaging has revealed DMN suppression by salient stimuli, the cellular mechanisms orchestrating this process remain unknown. Using whole-brain computational modeling informed by neuronal biophysics and retrograde tracer-derived directional mouse brain connectomics, we demonstrate that stimulation of the insula node of the salience network suppresses DMN activity, whereas cingulate cortex stimulation produces antagonistic effects, enhancing retrosplenial cortex activity. Prelimbic cortex stimulation showed intermediate patterns, partially replicating insula-mediated suppression while failing to suppress cingulate regions, suggesting its role as a functional bridge between networks. Systematic brain-wide analysis confirmed the insula’s unique pattern of simulated DMN suppression. Comprehensive parameter space exploration demonstrated that DMN emergence as a functionally segregated network is robust across wide ranges of excitatory-inhibitory balance regimes and cholinergic modulation. However, outside these boundaries, DMN integrity breaks down through three distinct failure modes: loss of responsiveness, reversal of suppression to enhancement, and network fragmentation. The retrosplenial cortex emerged as a particularly vulnerable regulatory hub whose excitatory-inhibitory disruption reversed normal suppression patterns across the DMN, while prelimbic cortex demonstrated remarkable robustness. Brain-wide analysis also identified a functionally segregated frontal network displaying antagonistic dynamics with the DMN. Our findings provide mechanistic insights into DMN robustness and vulnerability, establishing a framework that links cellular excitatory-inhibitory balance to large-scale network dynamics. This model could explain how region-specific disruptions can produce the heterogeneous patterns of DMN dysfunction observed across brain disorders.

**Significance Statement:** To respond to important external events, the brain must suppress internal thought processes implicating the default mode network. This suppression fails in psychiatric conditions, that also involve imbalances between excitatory and inhibitory neurons. However, the connection between cellular imbalances and default-mode-network dysfunction has remained unclear. We used brain-wide computer simulations incorporating neuronal properties to understand how imbalance at the cellular scale disrupts network function. Our simulations reveal the precise excitatory-inhibitory balance needed for normal suppression. Additionally, we identified distinct failure modes and discovered that certain brain hubs are more vulnerable than others to disruption. Our findings reveal how cellular alterations scale up to cause network-level dysfunction, potentially guiding treatments targeting specific brain regions in individual patients.

## Introduction

Cognition and its dysfunction in brain disorders emerges from how the brain dynamically regulates activity across distributed networks. The default mode network (DMN) has emerged as a key player in cognitive function, exhibiting strongly synchronized activity and characteristic suppression by salient external events ^1–4^. While extensive research has revealed the DMN’s crucial role in internal and self-referential mental processes including autobiographic memory and social cognition ^1,5,6^, the cellular and circuit mechanisms that enable its coordinated activation and suppression remain poorly understood. This mechanistic gap between cellular and network levels has profound implications for psychiatry and neurology, limiting our ability to develop targeted interventions for disorders characterized by network dysfunction.

Altered DMN function, particularly impaired suppression by salient external stimuli, has emerged as a robust biomarker across psychiatric disorders including autism and schizophrenia ^7–10^. DMN hyperactivity during tasks requiring external attention has been linked to impaired attention and cognitive flexibility ^11–14^, social deficits ^10^, and maladaptive rumination ^15^ across various conditions. Intriguingly, disorders with altered DMN function are hypothesized to involve excitation-inhibition (E/I) imbalance^16,17^, with converging evidence from human neuroimaging^10,18 19–21^ as well as animal models ^22,23^.

These parallel observations raise a fundamental question: could DMN function and dysfunction and E/I imbalance be causally linked? If so, how do cellular-level E/I disruptions shape neural circuit function to cause the network-level abnormalities observed in clinical populations? And critically, which specific DMN nodes are most vulnerable to such E/I disruptions in terms of their impact on global network dynamics? These questions have remained unanswered due to methodological challenges in bridging scales from neurons to large-scale brain networks, hampering the development of treatment strategies targeting the root causes of DMN dysfunction.

Recent technological advances have begun to address these challenges. Optogenetic manipulation combined with whole-brain imaging has enabled identification of a DMN homologue in rodents, anchored in cortical midline structures including retrosplenial cortex (RSC), cingulate cortex (Cg), and prelimbic cortex (PrL) ^4,24–26^. Critically, optogenetic studies demonstrated that DMN suppression can be induced by stimulating the anterior insula, a key hub of the salience/cingulo-opercular network ^4^. This finding provided causal evidence for mechanisms behind rodent DMN suppression, parallelling the homologous phenomenon observed in human neuroimaging studies. However, the cellular and circuit mechanisms, and specifically the role of E-I balance, underlying this network-level interaction remain unknown. Furthermore, the differential effects of stimulating various nodes within and outside the DMN have not been systematically compared, limiting understanding of specific roles of the insula versus DMN regions in network regulation.

Here, we address these critical knowledge gaps through biophysically informed computational modelling that bridges cellular mechanisms with large-scale network dynamics. We leverage The Virtual Brain (TVB) simulator ^27–31^ to implement novel mean-field models of neural dynamics ^32^. Our approach integrates the Allen Brain Atlas connectome, which provides experimentally validated, directional connectivity data based on viral tract tracing across 426 brain regions ^33^. This comprehensive dataset reveals the precise anatomical organization of circuits linking insula, DMN, and other brain regions, overcoming the directional ambiguity and limitations of crossing-fibers inherent in diffusion imaging ^34^. By incorporating this high-resolution connectome with biophysically informed excitatory and inhibitory neural dynamics, we create an analytic framework spanning from cellular properties to large-scale network interactions. This multi-scale approach enabled us to systematically investigate how cellular and circuit mechanisms give rise to DMN dynamics while maintaining biological plausibility and revealing principles of network organization.

Our study addresses six key interconnected objectives that systematically build understanding of DMN regulation mechanisms. First, we model DMN suppression by insula stimulation to identify the putative mechanisms that reproduce empirical findings ^4^. Second, we contrast the effects of insula stimulation against stimulation of Cg and PrL regions as well as sensory and motor regions in our model. Third, we examine how region-specific E/I imbalance alters DMN suppression patterns. Fourth, we analyze brain-wide responses to insula stimulation, focusing on regions beyond the DMN to characterize potential homologues of the human frontoparietal network. Fifth, we explore the parameter space governing E/I balance and DMN suppression, to identify parameter regimes where DMN suppression is maintained versus where it breaks down. Finally, we examine robustness of DMN emergence as a functionally segregated network across parameter space, providing mechanistic insights into network robustness and vulnerability.

The TVB model incorporates biophysical parameters of excitatory and inhibitory neurons (**Fig. 1**). Each brain region comprises interacting E and I neural populations whose dynamics are governed by Adaptive Exponential (AdEx) integrate-and-fire equations derived from single-neuron biophysics ^32^. These regional nodes are connected according to the empirically determined connectome from the Allen Mouse Brain Atlas, comprising 426 regions with directional connectivity weights based on viral tract-tracing ^33^. This architecture enables investigation of how cellular properties such as E/I balance shape large-scale network dynamics.

**Figure 1.**
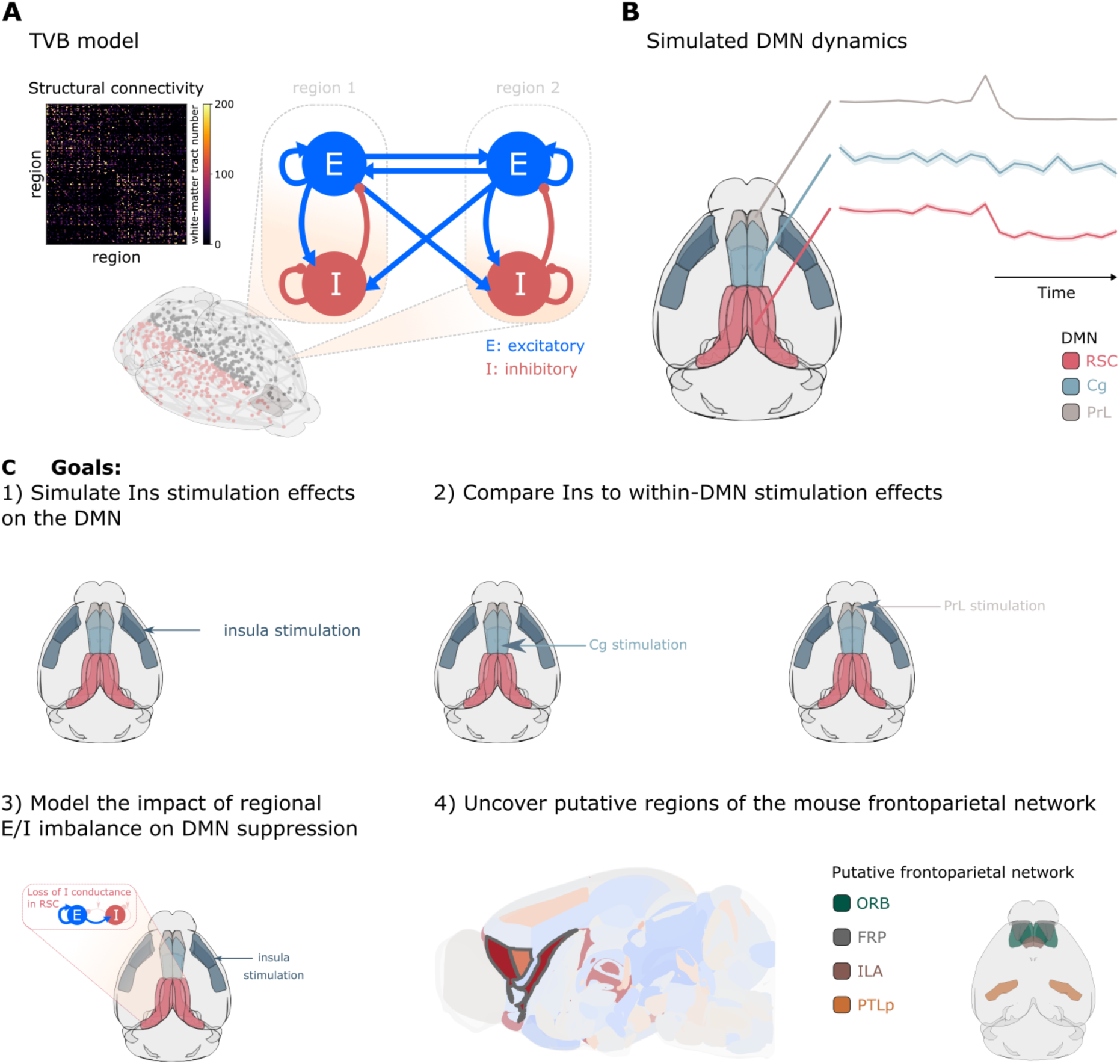
Model architecture and study objectives. **(A)** The Virtual Brain model architecture incorporating excitatory (E) and inhibitory (I) neuronal populations with reciprocal interactions. **(B)** Left: Anatomical delineation of default mode network (DMN) nodes including retrosplenial cortex (RSC), cingulate cortex (Cg), and prelimbic cortex (PrL). Right: Representative simulated population firing rate dynamics. **(C)** Schematic overview of study objectives.

Our approach reveals how E-I balance mechanisms could give rise to coordinated network-level phenomena previously observed in empirical studies but not mechanistically explained. Results identify specific circuit pathways potentially mediating DMN suppression and demonstrate how localized E/I disruptions can cause distinct patterns of network dysfunction. Our findings reveal how E-I balance alterations propagate through circuits to produce the network-level dysfunction.. By linking specific regional E/I disruptions to distinct patterns of altered network function, our computational approach could inform precise diagnostic approaches and guide development of targeted interventions that address specific circuit abnormalities.

## Results

### TVB model accounts for DMN suppression upon insula activation

We investigated whether biophysically informed neural dynamics coupled with empirical connectomes can reproduce this phenomenon and reveal its underlying mechanisms. We implemented a whole-brain model comprising 426 regional nodes, each containing E and I neural populations whose dynamics are governed by mean-field integrate-and-fire equations (see Methods). Regions were connected according to the Allen Mouse Brain Atlas connectome derived from viral tract-tracing experiments ^33^. To test whether our model could reproduce DMN suppression as observed in optogenetic experiments, we simulated periodic stimulation of the right insula (0.5 seconds every 2 seconds) and analyzed the resulting changes in DMN activity across 100 independent model realizations (**Fig. 2A**).

**Figure 2.**
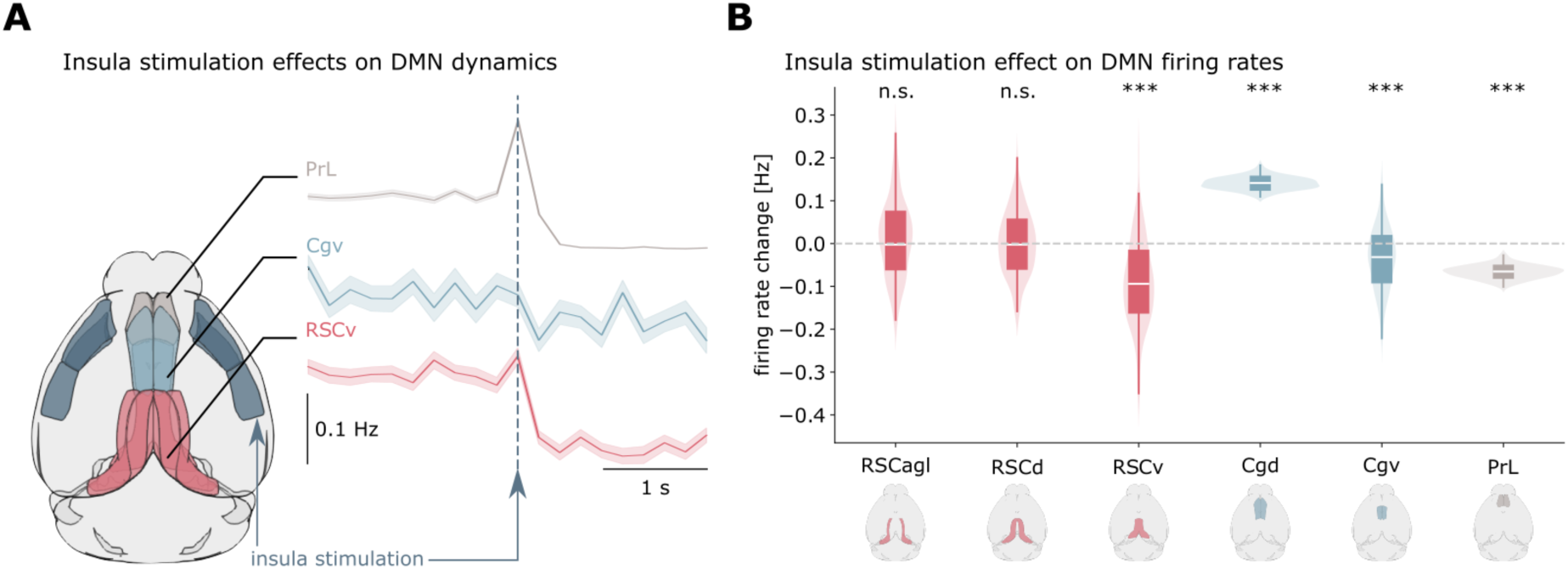
Insula stimulation produces coordinated DMN suppression. (**A)** Schematic of insula stimulation. Time courses show suppression of ventral RSC (RSCv), ventral Cg (Cgv), and PrL following insula stimulation. **(B)** Violin plots quantifying firing rate changes across DMN subregions, averaged over 100 model realizations. Results demonstrate coordinated suppression of RSCv, Cgv, and PrL, with enhanced activation of dorsal Cg (Cgd), revealing functional heterogeneity within the DMN. These findings recapitulate empirically observed DMN suppression by insula activation.

Our simulations demonstrated significant suppression of key DMN nodes following insula stimulation. Specifically, the ventral retrosplenial cortex (RSCv), ventral cingulate cortex (Cgv), and prelimbic cortex (PrL) all showed reduced activity (Wilcoxon test, p < 0.001, FDR corrected; **Fig. 2B**). Interestingly, we observed functional heterogeneity within the cingulate cortex, with dorsal Cg (Cgd) exhibiting enhanced rather than suppressed activity following insula stimulation (*p* < 0.001, FDR corrected). Parameter exploration revealed that DMN suppression emerged robustly when E-to-I connections between regions were sufficiently strong relative to E-to-E connections (ratio between 1.2 and 2.8).

Our model successfully reproduces the empirically observed pattern of DMN suppression following insula stimulation, providing a mechanistic explanation for this phenomenon. The results emphasize that E-to-I connectivity across brain regions is a necessary circuit-level mechanism for us to model DMN suppression, enabling inhibitory neurons to silence local excitatory activity upon receiving inputs from the insula.

### Stimulation of prefrontal DMN nodes produces effects distinct from insula stimulation

While insula stimulation suppresses DMN activity, it remains unknown to what extent suppressive effects are specific to the insula. Specifically, the differential effects of stimulating nodes within the DMN versus the insula, a salience network hub with antagonistic DMN relationships, have not been systematically examined. Such comparisons can reveal the causal relationship between these networks and clarify the functional roles of different DMN nodes. To address this, we compared the effects of stimulating two putative DMN regions— Cg and PrL— with our established insula stimulation paradigm. This allows us to directly compare the effects of within-DMN to those of insula stimulation on the RSC, believed to be a structural ^35^ and functional ^24,26^ hub anchoring the rodent DMN. Using identical stimulation parameters across targets, we analyzed the resulting activity patterns in DMN subregions, focusing on the RSC subdivisions, ventral and dorsal cingulate (Cgv, Cgd), and PrL. We quantified differences in stimulation effects using Wilcoxon tests with FDR correction (**Fig. 3**).

**Figure 3.**
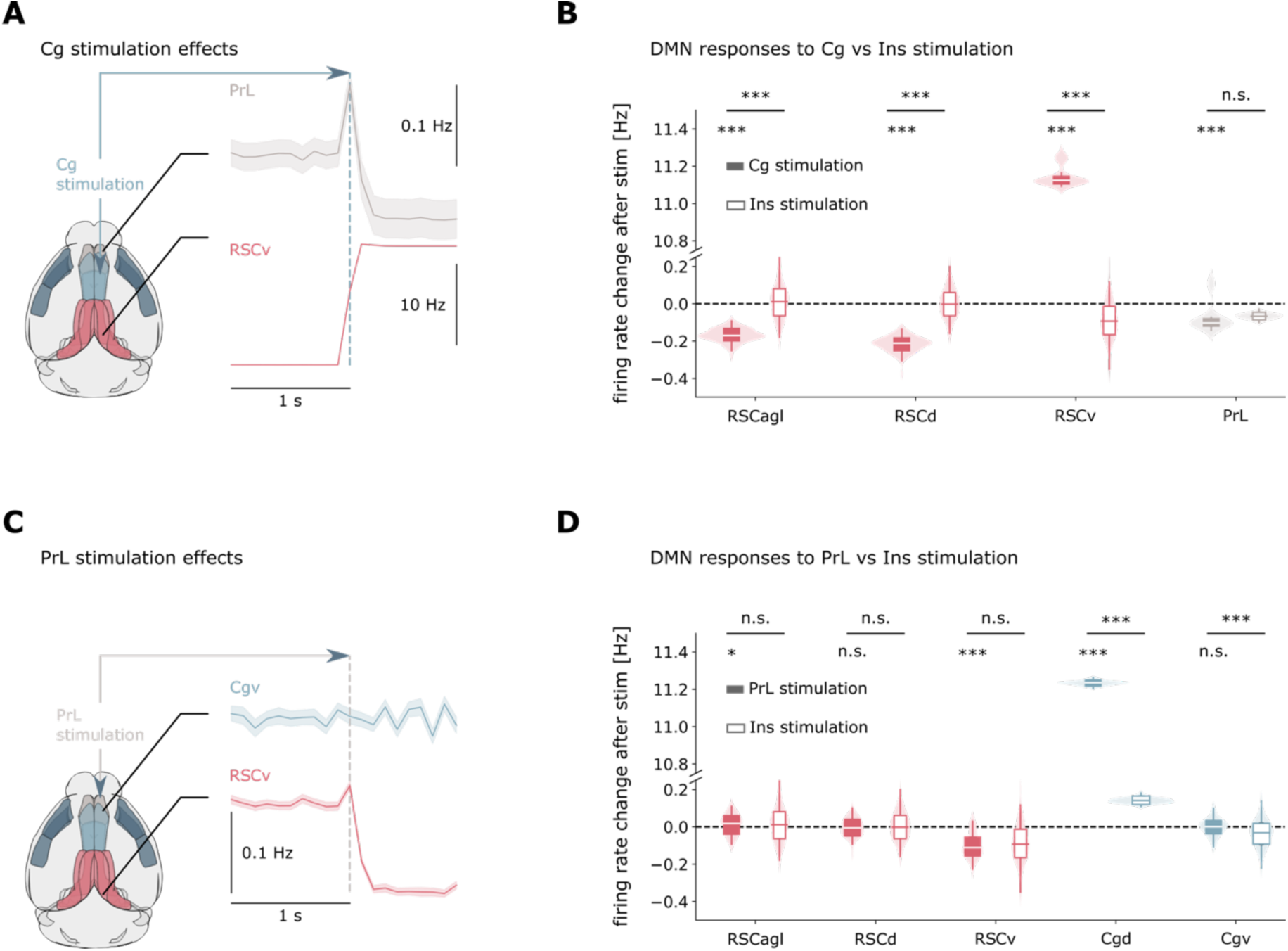
Distinct network effects of DMN node stimulation reveal hierarchical control mechanisms. **(A-B)** Cingulate cortex stimulation produces effects opposing those of insula stimulation on RSC activity, demonstrating network antagonism between DMN and salience network circuits. **(C-D)** Prelimbic cortex stimulation partially recapitulates insula effects on RSC but produces distinct effects on Cg, suggesting PrL may serve as an interface region between networks. Results reveal hierarchical organization of inter-network control mechanisms.

Cingulate stimulation produced strikingly different effects compared to insula stimulation. While insula activation suppressed the RSCv, Cg stimulation significantly enhanced RSCv activity and suppressed the PrL (*p* < 0.001). Conversely, Cg stimulation suppressed agranular and dorsal RSC (RSCagl, RSCd), regions that were less affected by insula stimulation. Effects of stimulating either the Cgv or the Cgd cingulate subdivisions separately yielded similar findings (**Fig. S2A-B**). Effects of Cg stimulation on the RSC were reciprocal, as RSC stimulation (**Fig. S2C-F**) enhanced the Cg overall while also suppressing the PrL (*p* < 0.001). PrL stimulation showed a more complex pattern: like insula stimulation, it suppressed the RSCv, producing effects that were significantly different from those of insula on the RSC (*p* > 0.05). However, unlike insula stimulation, PrL activation failed to suppress the Cgv, indicating a partial dissociation in network effects.

These results suggest a causal antagonism between the DMN and the insula, where cingulate stimulation produces effects opposite to insula stimulation on key DMN nodes in simulations. The distinct pattern observed with PrL stimulation suggests this region may appear as an interface between networks, sharing characteristics with both DMN and salience circuits.

### Insula stimulation produces unique distributed DMN suppression distinct from sensory and motor cortices

To test the specificity of insula-mediated DMN suppression, we systematically compared effects of insula stimulation against stimulation of other brain regions. This analysis revealed that the insula produces a distinctive pattern of distributed DMN suppression fundamentally different from simulations of other brain regions (**Fig. S3**). While the insula produces coordinated suppression across multiple DMN subdivisions (RSCv, Cgv, PrL), sensory and motor regions generate either selective suppression of individual nodes or mixed patterns combining suppression and enhancement across different subdivisions. For example, visual cortex stimulation suppresses Cgv and PrL but enhances RSCv, opposite to insula effects on RSCv (Fig. S3).

### Regional E/I balance controls DMN suppression patterns

E/I imbalance has been implicated in various psychiatric disorders, including autism ^16,17^, which also show altered DMN function. However, the causal relationship between cellular E/I balance and network-level DMN function remains unknown. To address this, we systematically investigated how localized E/I imbalance affects simulated DMN suppression by selectively reducing inhibitory synaptic conductance in different DMN regions. We manipulated I synaptic conductance model parameters (from baseline 5 nS to 4 nS) controlling I synaptic strength and hence E/I balance in RSC, Cg, and PrL separately, while measuring the resulting changes in network dynamics during insula stimulation (**Fig. 4**). Statistical comparisons were performed using Wilcoxon tests with FDR correction.

**Figure 4.**
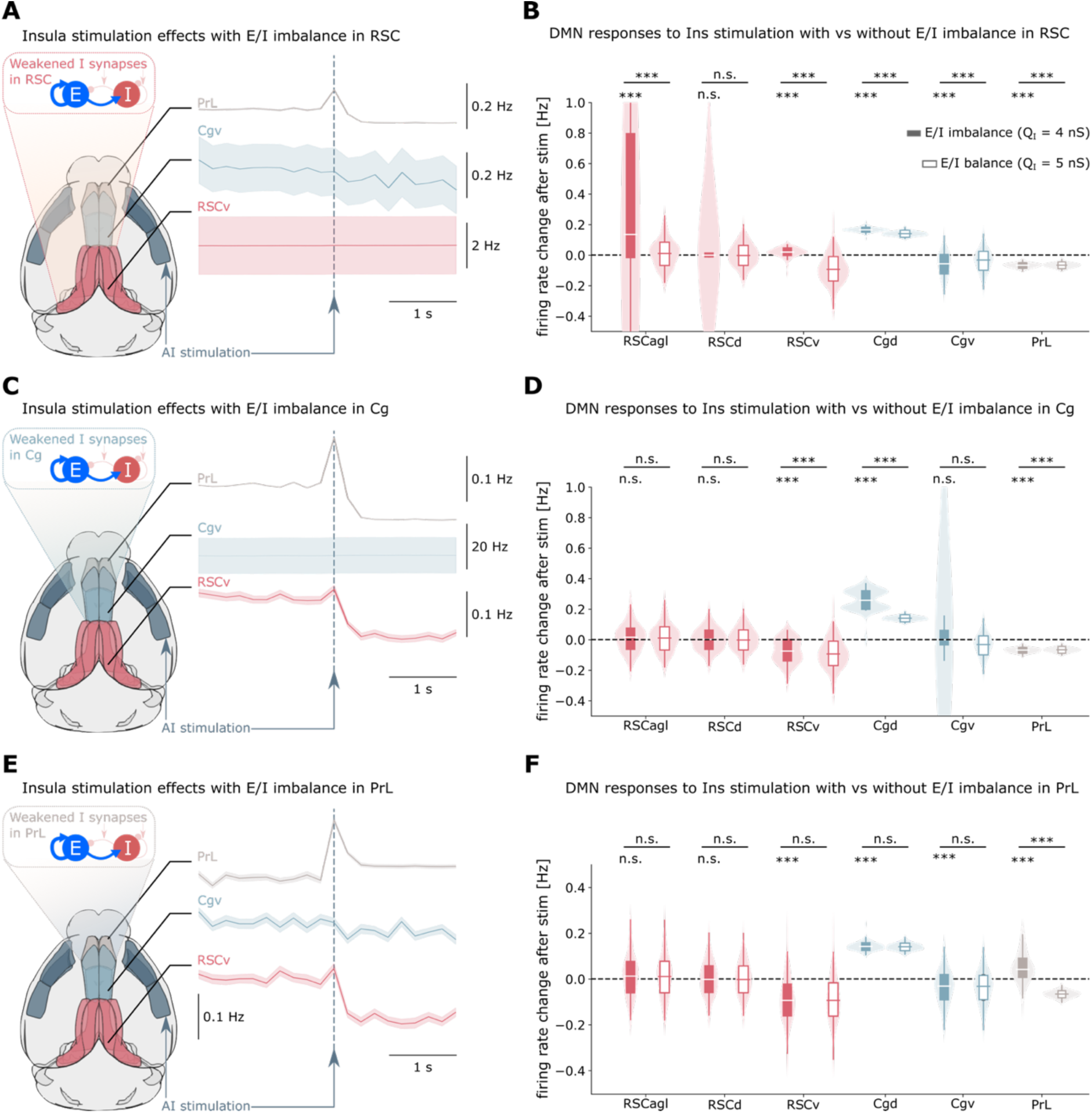
Regional E/I imbalance disrupts normal DMN suppression patterns. **(A-B)** E/I imbalance in RSC reverses the normal suppression pattern: both dorsal and ventral RSC show enhanced activity following insula stimulation, while suppression is paradoxically increased in Cgv and PrL. Regional E/I imbalance was modeled by weakening inhibitory synaptic conductance. **(C-D)** E/I imbalance in Cg abolishes Cgv suppression, attenuates RSCv suppression, and enhances PrL suppression, suggesting multiple pathways mediate DMN suppression. **(E-F)** E/I imbalance in PrL enhances local activity without affecting suppression in RSC or Cg, consistent with PrL’s relative disengagement from the DMN and potential dual involvement in both DMN and salience network circuits. Results demonstrate how localized cellular perturbations can produce widespread network-level alterations.

**Figure 5.**
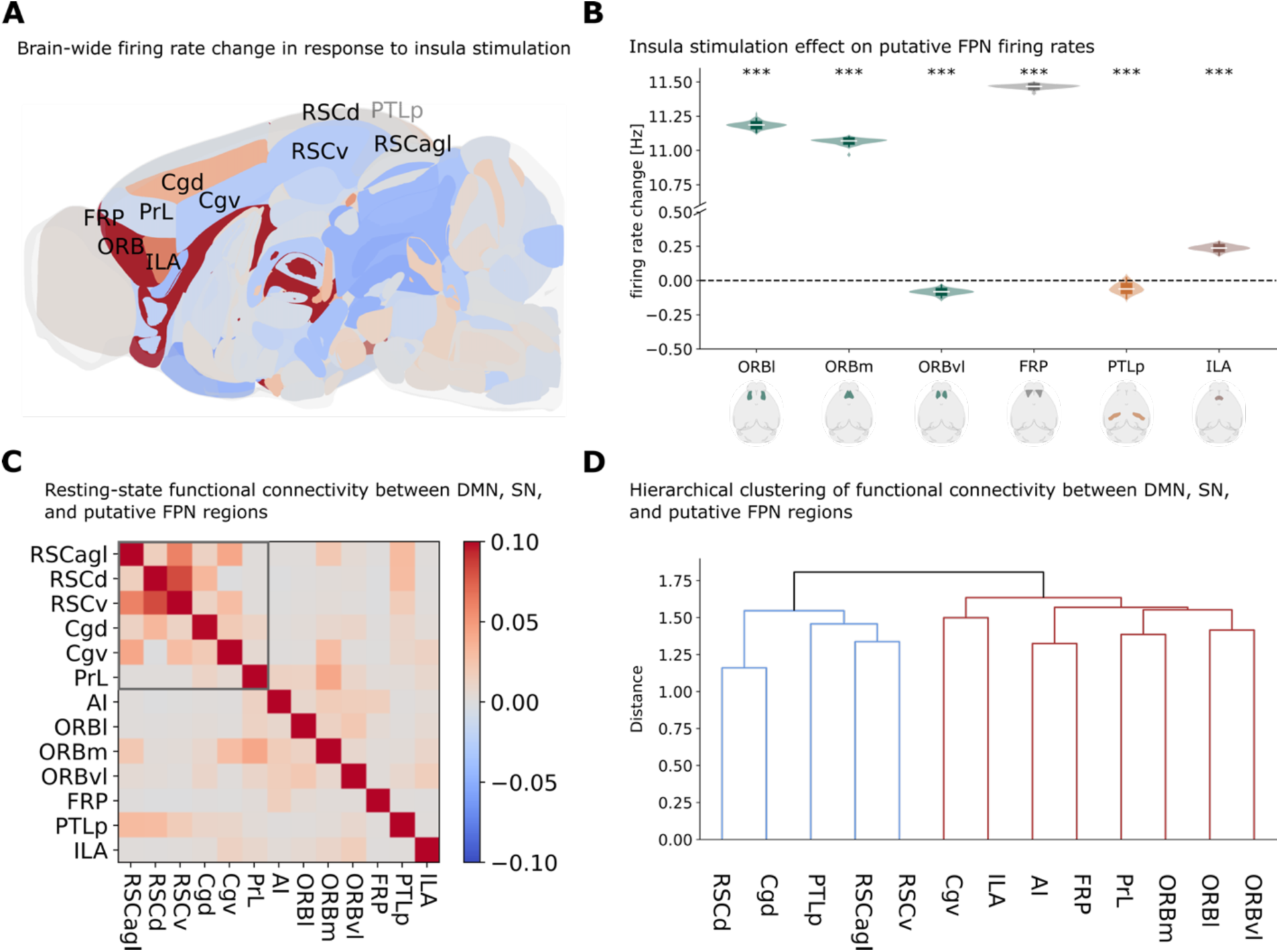
Identification and characterization of a putative rodent frontal network. **(A)** Sagittal view of simulated right-hemisphere firing rate responses to right insula stimulation. Colors indicate firing rate changes (red: enhancement; blue: suppression). Visible (black) and hidden (gray) regions of the DMN, salience network, and putative frontal network are labeled. **(B)** Violin plots quantifying firing rate changes across putative frontal network subregions, demonstrating selective suppression of orbital cortex lateral (ORBl), orbital cortex medial (ORBm), frontal pole (FRP), and caudoputamen (CP). **(C-D)** Resting-state functional connectivity matrix (C) and hierarchical clustering dendrogram (D) among right-hemisphere DMN, salience network, and putative frontal network subdivisions. Hierarchical clustering reveals a spontaneous cluster of DMN regions (D, red) that includes posterior parietal cortex (PTLp) but excludes PrL, which aligns with salience and frontal networks during resting-state but shifts toward DMN during insula stimulation.

Each regional E/I disruption produced a distinct pattern of altered DMN suppression. E/I imbalance in the RSC caused a complete loss of insula-mediated suppression in RSC subregions, effectively reversing the normal response pattern, while paradoxically increasing suppression in Cgv and PrL (**Fig. 4A-B**). E/I disruption in Cg led to loss of Cgv suppression, attenuated RSCv suppression, and enhanced PrL suppression (**Fig. 4C-D**). When E/I imbalance was introduced in the PrL, suppression was abolished specifically in this region while RSC and Cg responses remained largely intact (**Fig. 4E-F**). These region-specific patterns were maintained across different levels of E/I imbalance: consistent findings were found at a less severe E/I imbalance, arising from I conductance of 4.5 nS (**Fig. S2**). This result underscores the robustness of our findings to the value of this biophysical parameter.

These results reveal that sufficient inhibitory conductance is necessary to simulate normal DMN suppression, and that the network-wide consequences of E/I disruption could depend critically on which region is affected.

### TVB modeling reveals a putative mouse homolog of a frontal network

The frontoparietal network plays a critical role in cognitive control and working memory in humans and is activated during salient stimulus processing. However, whether a homologous network exists in rodents remains unclear. We analyzed brain-wide responses to insula stimulation, focusing on regions beyond the established DMN and insula. We examined activity changes in frontal regions including the orbitofrontal cortex (ORB), frontal pole (FRP), and infralimbic cortex (ILA), as well as the posterior parietal cortex (PTLp), which might constitute a putative mouse frontoparietal network (**Fig. 6A-B**).

**Figure 6.**
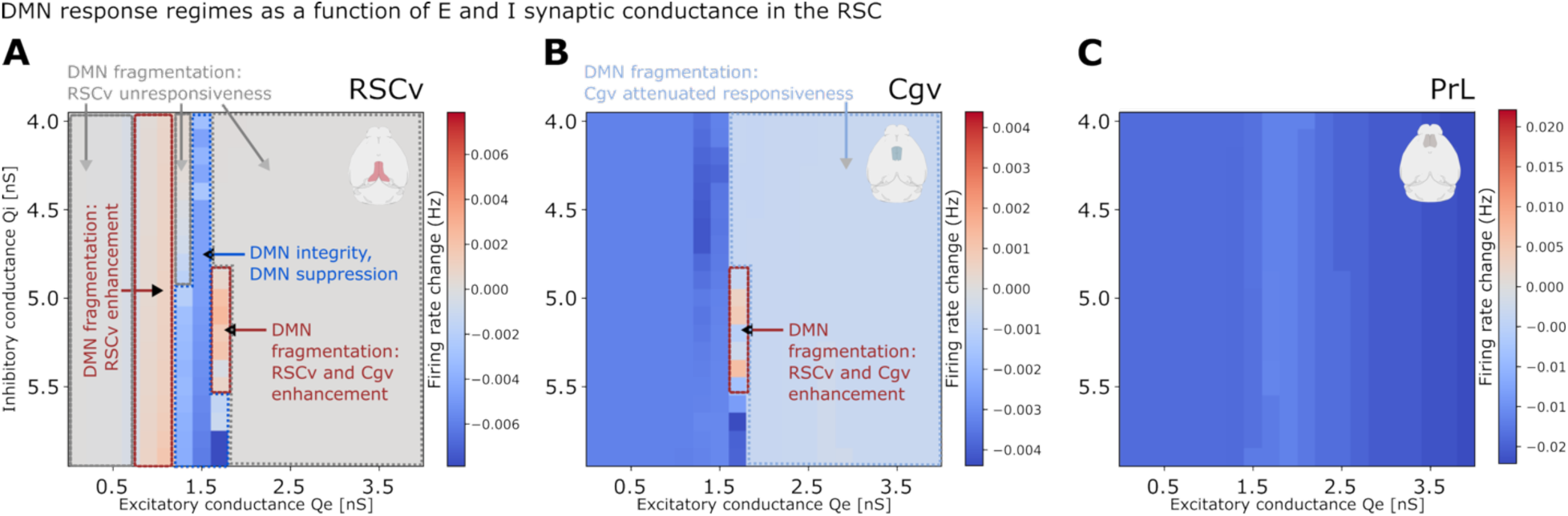
Parameter space exploration reveals multiple failure modes of DMN suppression. Heatmaps show simulated firing rate responses to insula stimulation in **(A)** RSCv, **(B)** Cgv, and **(C)** PrL as a function of excitatory and inhibitory synaptic conductance in RSC. Simulations used fixed parameters: E-to-I/E-to-E conductance ratio = 1.4 and spike-frequency adaptation *=* 0 nS. Results demonstrate the existence of parameter regimes supporting robust DMN suppression and integrity, as well as regimes associated with distinct DMN suppression failure modes, accompanied by DMN fragmentation: local unresponsiveness and attenuation, local reversal from suppression to enhancement, and distributed reversal to enhancement across multiple DMN regions.

Insula stimulation produced robust activation in specific frontal regions, with lateral (ORBl) and medial (ORBm) orbitofrontal cortex, frontal pole, and caudoputamen all showing significant increases in firing rates (*p* < 0.001, FDR corrected). This activation pattern showed regional specificity; ventrolateral orbitofrontal cortex (ORBvl) and posterior parietal cortex were suppressed rather than enhanced (**Fig. 6B**).

Next, we characterized the intrinsic network organization of these regions in relation to DMN and the insula (**Fig. 6C-D**). Functional connectivity analysis revealed that orbitofrontal regions showed strong coupling with insula subdivisions, while the PTLp was more strongly correlated with the RSC. Hierarchical clustering identified two distinct network clusters: a DMN-like cluster containing RSC, Cgd, and PTLp, and a frontal cluster containing ORB, FRP, PrL, ILA grouped with the AI (**Fig. 6D**).

These results identify a functionally segregated putative frontal network in mice comprising orbitofrontal cortex and frontal pole, and infralimbic regions that shows distinct dynamics from the DMN. These findings generate hypotheses about a mouse homolog of the human frontal network, though without a lateral posterior parietal component, that could be tested in neuroimaging experiments in behaving animals.

### Comprehensive parameter space analysis reveals regimes of DMN integrity and suppression

To identify conditions under which fundamental DMN properties are maintained versus where they break down, we systematically explored how local E/I balance in RSC, inter-regional coupling, and cholinergic neuromodulation interact to shape DMN responses to insula stimulation. This comprehensive analysis spanning over 14,000 simulations across multiple parameter dimensions revealed specific regimes supporting robust DMN organization and suppression, as well as distinct modes of breakdown (Fig. 6-7, S4-S8, Tables S1, S2).

**Figure 7.**
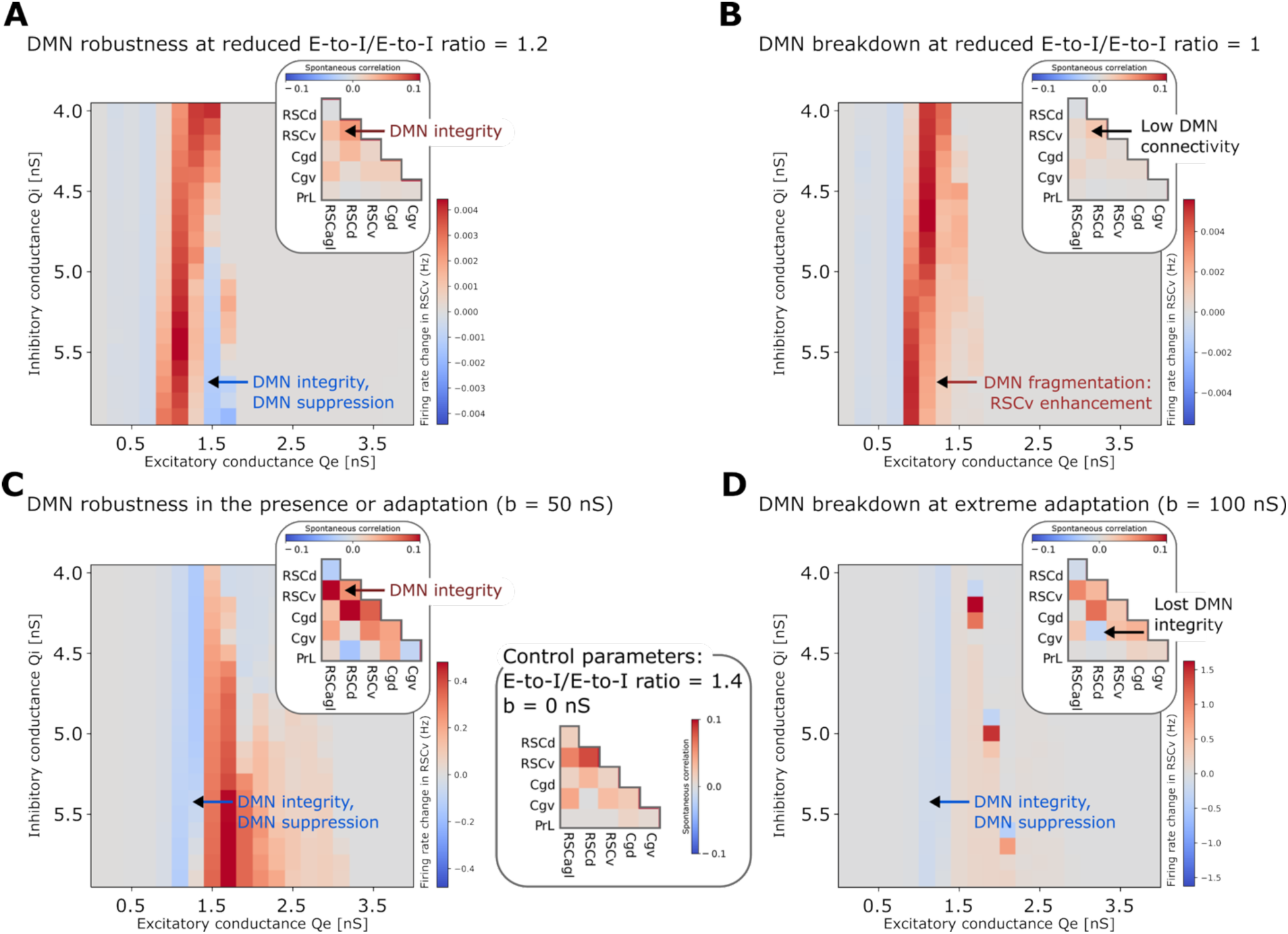
Global circuit parameters differentially affect DMN suppression and network integrity. Heatmaps show simulated RSCv responses to insula stimulation as a function of excitatory and inhibitory synaptic conductance in RSC. **(A-B)** Effects of varying E-to-I/E-to-E conductance ratio at 1.2 (A) and 1.0 (B) with zero adaptation. **(C-D)** Effects of modeled cholinergic neuromodulation via spike-frequency adaptation at 50 nS (C) and 100 nS (D) with fixed E-to-I/E-to-E ratio of 1.4. Insets show within-DMN functional connectivity during spontaneous activity (no stimulation) for each parameter set, with control parameters (E-to-I/E- to-E ratio = 1.4, adaptation = 0 nS) shown in panel C, right. Results demonstrate robustness of DMN emergence as a coherent network: reduced E-to-I/E-to-E ratio causes DMN response fragmentation and reversal but not complete breakdown of DMN emergence, while increased adaptation disrupts DMN coherence by causing fragmentation into multiple spontaneously emerging subnetworks without preventing suppression.

DMN emergence as a functionally segregated network, with insula-mediated suppression, was robust across both local and global model parameters: *Q*_*E*_ between 0.7-1.7 nS and *Q*_*I*_ between 4.5-5.5 nS in the RSC (Fig. 6, Fig S4 for other regional subdivisions), inter-regional E-to-I/E-to-E ratios between greater than 1, and adaptation below 100 nS (Fig. 7A-C, Fig. S5-S6). Within these boundaries, RSCv, Cgv, and PrL showed consistent suppression following insula stimulation, with PrL demonstrating particularly robust suppression across nearly all parameter combinations.

Outside these regimes, DMN integrity broke down through distinct modes (Fig. 6, Table S1). First, localized loss of responsiveness occurred when *Q*_*E*_ in the RSCv was too low (<0.7 nS) or too high (>1.7 nS). In these regimes, the RSCv showed minimal firing rate changes in response to insula stimulation. Second, localized reversal from suppression to enhancement of the RSCv emerged when *Q*_*E*_ in RSC was moderate (0.7-1.3 nS), combined with a low inter-regional E-to-I/E-to-E coupling (ratio = 1) (Fig. 7B). Third, distributed reversal from suppression to enhancement occurred when E/I balance strongly favored excitation, specifically at high *Q*_*E*_ (> 1.6 nS) combined with low *Q*_*I*_ (5-5.5 nS) in the RSC. This condition produced enhancement of both posterior and anterior DMN nodes (RSCv and Cgv) following insula stimulation (Fig. 6A-B). Notably, PrL suppression remained robust across all parameter perturbations tested, demonstrating that different DMN nodes show markedly different vulnerabilities to E/I disruptions (Fig. 6C). Fourth, our model also identified an additional network fragmentation under extreme neuromodulatory conditions. At extreme adaptation levels (b=100 nS), representing states of minimal cholinergic tone, the DMN split into ventral (RSCagl, RSCv, Cgv) and dorsal (RSCd, Cgd, PTLp) functional subclusters (Fig. 7D, Fig. S8). In contrast, reduced inter-regional E-to-I/E-to-E coupling (ratio = 1) weakened within-DMN functional connectivity (Fig. 7B) but did not disrupt DMN emergence as a cluster functionally segregated from the insula and frontal regions (Fig. S8).

These results suggest that DMN emergence as a functionally segregated network is robustly preserved across wide parameters ranges and identifies four distinct failure modes when E/I balance and neuromodulatory adaptation are perturbed. RSC emerged as particularly vulnerable to E/I balance disruptions, while PrL demonstrated stability across nearly all perturbations (**Table S2**).

## Discussion

Our computational investigation reveals fundamental mechanisms linking cellular-level E-I balance to large-scale brain network dynamics, addressing critical gaps in understanding how the DMN is regulated. Using biophysically informed whole-brain modeling, we demonstrate how specific cellular and circuit properties give rise to coordinated DMN function and its modulation by the insula. Our analysis yields several key insights bridging microscale and macroscale brain organization. First, our model predicts that DMN suppression by insula stimulation requires specific excitatory-inhibitory circuit interactions, particularly a balanced ratio of E-to-I versus E- to-E connectivity between regions. Second, insula stimulation produces coordinated DMN modulation that differs fundamentally from Cg and PrL stimulation, with antagonistic effects on key nodes like the RSC.

Third, our simulations demonstrate how localized disruptions in E/I balance ^16,17^ could propagate to cause distinct patterns of network dysfunction, with RSC emerging as a particularly vulnerable node whose disruption affects DMN-wide suppression patterns. Fourth, systematic parameter space exploration revealed the specific regimes where DMN suppression is maintained versus where it breaks down, identifying non-linear interactions between local synaptic conductance, inter-regional coupling, and cholinergic modulation. Fifth, we uncovered evidence for a segregated frontal network displaying opposing responses to the DMN. Finally, comprehensive parameter exploration demonstrated remarkable robustness of DMN emergence as a functionally segregated network across physiologically relevant ranges of E/I balance and neuromodulation, while revealing specific vulnerabilities where DMN organization and insula-induced suppression break down under extreme parameter combinations. These findings provide a computational framework linking E-I balance cellular mechanisms with large-scale function of the DMN in tasks, bridging traditionally separate domains of neuroscience investigation ^32^, with relevance for understanding brain function and its disruption in disorders.

### Cellular mechanisms underlying insula-mediated DMN suppression

Our biophysically informed model successfully reproduces empirically observed DMN suppression by insula stimulation, providing mechanistic insights into this fundamental network interaction. Specifically, our simulations predict that insula stimulation leads to suppression of key DMN nodes, including ventral retrosplenial cortex (RSCv), ventral cingulate cortex (Cgv), and prelimbic cortex (PrL), consistent with experimental observations in rodents ^2,36^.

Our analysis revealed that E-to-I connectivity across regions is a crucial mechanism enabling DMN suppression in our model. Extensive parameter space exploration demonstrated that DMN suppression only emerges when E-to-I connections between regions are sufficiently strong relative to E-to-E connections, specifically when this ratio falls between 1.2 and 2.8. This finding suggests a mechanistic hypothesis: insula stimulation preferentially activates inhibitory interneurons in DMN regions, which suppress local excitatory activity. While human neuroimaging studies have established correlational relationships between insula and DMN activity ^3^, and optogenetic studies have demonstrated causality ^4^, our model provides the first mechanistic investigation into how this interaction may occur at a cell-type-specific resolution level.

Importantly, these findings cannot be straightforwardly understood from naïve structural connectivity analyses alone. In the Allen mouse connectome atlas, the insula has no direct monosynaptic connections to either RSC or Cg ^33,35^. Our simulations reveal that DMN suppression emerges from indirect pathways combined with regional E/I balance mechanisms, insights not readily apparent from structural connectivity alone. The model generates specific predictions about the spatial specificity and temporal dynamics of suppression across DMN nodes.

### Network control: Differential effects of insula versus DMN node stimulation

Comparison between stimulation of insula and DMN nodes provides crucial insights into network control principles. Our results demonstrate that insula stimulation can produce coordinated DMN suppression that differs fundamentally from stimulation of Cg and PrL regions in our simulations. Specifically, stimulation of Cgd showed effects opposite to insula stimulation, enhancing rather than suppressing RSC subdivisions and providing evidence for causal antagonism between specific DMN and salience network circuits ^1–4^. Stimulation of Cgv produced a mixed pattern with partial suppression of RSCd but enhanced activity in RSCv, suggesting functional heterogeneity within the cingulate cortex itself. These anatomical dissociations align with experimental evidence showing distinct connectivity patterns and functional roles of dorsal versus ventral cingulate regions in rodents ^33^ and humans ^37^.

In contrast, PrL stimulation partly mimics insula effects on RSC, with no significant differences between PrL and insula stimulation on RSC activity. Like the insula, PrL stimulation enhanced Cgd activation, but unlike the insula, failed to suppress Cgv. This intermediate pattern suggests the PrL’s functional properties bridge DMN and salience networks. This functional position is supported by the PrL’s unique anatomical organization, as it receives direct inputs from the insula unlike other DMN regions ^33^. Hierarchical clustering analyses of GCaMP recordings further support this organization, showing RSC and Cg closely aligned while PrL is more distant^25^.

These results demonstrate causal antagonism between the DMN and task-positive networks, where dorsal cingulate stimulation produces effects opposite to insula stimulation on key DMN nodes. This functional organization helps explain how the brain maintains separation between internally-oriented (DMN) and externally-oriented (salience network) attentional states while allowing controlled transitions between them. The reciprocal relationship between Cgd and insula effects provides a mechanistic basis for the antagonistic relationship observed between these networks during cognitive tasks, with implications for understanding disorders characterized by altered network switching ^3,38^.

Based on these findings, we predict that the insula serves as a primary control node driving coordinated suppression across specific DMN nodes, while nodes within and adjacent to the DMN provide more localized control. We further predict that the PrL’s ability to partially replicate insula effects while maintaining unique influence over Cg suggests it serves as a secondary control point for fine-tuning network interactions.

Our results also generate testable predictions about how stimulation of cingulate, prelimbic, sensory, and motor cortices differentially modulate DMN dynamics, predictions that can be validated through optogenetic experiments.

### E/I balance in network function and dysfunction

Our systematic modeling of E/I balance reveals its fundamental role in maintaining DMN function and provides a putative mechanistic link between cellular disruptions and network-level abnormalities. By methodically reducing inhibitory conductance in specific DMN nodes while measuring responses to insula stimulation, we identified regional vulnerabilities and circuit pathways through which cellular disruptions propagate.

Importantly, each region’s E/I disruption produced distinct patterns of network alterations. These region-specific patterns of altered network function have potential relevance for neuropsychiatric disorders. The heterogeneous E/I-induced changes we observed parallel complex patterns of altered connectivity documented in conditions like autism, where both hyper- and hypo-connectivity have been reported in different subdivisions of posterior cingulate and RSC ^39^. Our model suggests such mixed connectivity patterns may arise from localized E/I disruptions in specific network nodes.

Our findings provide testable predictions: reducing GABAergic inhibition specifically in RSC can reverse RSC and Cg suppression to enhancement but have minimal effects on the PrL. These predictions could be tested through cell-type-specific optogenetic or chemogenetic manipulations in mouse models.

### Distinct functional segregation of the DMN from frontal networks in the mouse brain

Understanding large-scale brain network organization in rodents is fundamental for translational neuroscience, yet substantial questions remain about homology to human networks. While recent studies have advanced understanding of the DMN and salience network in rodents ^24 4,40,41^, a clear rodent homolog of the human frontoparietal network has remained elusive. Our insula stimulation approach provides a novel model-based method to probe network interactions, addressing how the DMN may functionally relate to other large-scale networks in mice.

Our simulations reveal a functional cluster containing AI, PrL, ILA, Cgv, FRP, and orbitofrontal regions (ORBm, ORBl, ORBvl). The functional properties of these regions suggest they form a coherent network involved in executive control, decision-making, and salience processing— functions analogous to those performed by frontoparietal and salience networks in humans. Notably, mouse posterior parietal cortex (PTLp) aligns functionally with the DMN rather than with frontal regions, as demonstrated by both insula stimulation responses and intrinsic connectivity patterns in our simulations. This is consistent with the rodent PTLp’s structural links and navigation-related functional similarity with the RSC ^42^. These findings also generate testable predictions for experimental validation, including pertinent recording sites for examining frontal network engagement during cognitive control tasks in rodents.

### Robustness of DMN emergence and parameter regimes governing network integrity and breakdown

Our comprehensive parameter space exploration reveals a fundamental principle: DMN emergence as a functionally segregated network is remarkably robust, yet specific features of its dynamics, particularly insula-mediated suppression, depend on precise parameter regimes. This dissociation between network identity and network dynamics has important implications for understanding both normal function and pathological states.

DMN functional segregation from frontal and insular networks persisted across wide ranges of local E/I balance, inter-regional coupling ratios, and cholinergic modulation levels (**Fig. 6-7**). This robustness suggests that DMN organization reflects a fundamental architectural principle of brain organization, emerging from the convergence of anatomical connectivity patterns and cellular properties. The preservation of DMN identity even under substantial parameter perturbations may explain why this network is consistently identified across neuroimaging studies despite individual differences in underlying cellular and circuit properties.

Our analysis also identified distinct failure modes involving altered insula-mediated suppression and network fragmentation, which provided a novel framework for understanding heterogeneity in psychiatric disorders involving DMN dysfunction. Different individuals with autism or schizophrenia may show different failure modes depending on the specific nature and location of their E/I imbalances. For example, patients showing consistent DMN hyperactivation across contexts may have E/I imbalance favoring excitation (reversal mode), while those showing variable or absent DMN responses may have parameters near boundaries of responsiveness (loss of responsiveness mode). Comparing these computational failure modes to DMN activity patterns in mouse models of autism and schizophrenia could test whether specific genetic disruptions produce specific failure modes, generating mechanistic hypotheses about genotype-phenotype relationships ^43–45^. Additionally, our findings also provide testable predictions for optogenetic or chemogenetic experiments modulating E and I neurons selectively, as well as for pharmacological manipulations of spike-frequency adaptation via cholinergic neuromodulation.

### Limitations

Our mean-field modeling approach treats all excitatory neurons as identical and all inhibitory neurons as identical within each region, omitting complex intra-regional microcircuitry including cortical layers, columns, and distinct inhibitory subtypes (parvalbumin-positive, somatostatin-positive, VIP-positive interneurons). While information about cell-type-specific connectivity is emerging for some regions, it is not yet comprehensively available across the 426 regions of the Allen mouse brain connectome ^46^. Additionally, we modeled only excitatory pyramidal neurons and inhibitory interneurons, using the same E-I framework for cortical and subcortical structures. Cell types specific to subcortical structures have distinct biophysical properties not captured by our cortical neuron models. We incorporated only acetylcholine (modeled through spike-frequency adaptation), omitting dopamine, noradrenaline, and serotonin systems that modulate DMN function and are implicated in psychiatric disorders. Future work incorporating region-specific additional cell-types and neuromodulation systems could refine these predictions and reveal additional mechanistic insights into network function and dysfunction.

## Conclusions

Our study reveals how cellular-scale E/I balance mechanisms could orchestrate DMN dynamics, bridging a critical gap between molecular mechanisms and network-level phenomena. Through mean-field models incorporating tracer-derived directional connectomes, we identify putative circuit interactions underlying DMN suppression and demonstrate the anterior insula’s specialized role in coordinating distributed network regulation. Systematic comparison across all brain regions supports the insula’s unique profile of coordinated DMN suppression.

Comprehensive parameter space exploration revealed the precise conditions under which modeled DMN function is maintained versus where it breaks down through three distinct modes: loss of responsiveness, reversal to enhancement, and network fragmentation. Our simulations demonstrate that appropriate E/I balance is essential for normal DMN function, and that localized E/I disruptions propagate through specific circuit pathways to cause distinct patterns of network dysfunction. Critically, different DMN regions show differential vulnerability to E/I perturbations, with retrosplenial cortex emerging as a particularly sensitive regulatory hub whose disruption reverses normal suppression patterns.

Our computational framework links specific cellular perturbations like E/I imbalance and neuromodulation to network-level dysfunctions observed in psychiatric disorders, potentially informing precision diagnostic approaches and therapeutic strategies targeting region-specific circuit abnormalities. These mechanistic insights have implications for understanding both healthy brain function and developing targeted interventions for neuropsychiatric disorders characterized by network-level dysfunction.

## Methods

### Biophysical neural network model

To model the biophysical mechanisms underlying whole-brain dynamics and DMN function, we developed a biophysical model of adaptive exponential (AdEx) neurons of whole-brain dynamics in the mouse brain. The model was implemented in The Virtual Brain (TVB) simulator ^27,28^ based our previous implementation for the human brain ^32^. Each region comprises an E and an I neural populations whose firing rate dynamics reproduce the behavior of AdEx spiking neuron networks (Gerstner and Brette 2009). In these spiking network models, the neural membrane voltage *V* for a neuron of type *μ* ∈ {*E*, *I*} is described by

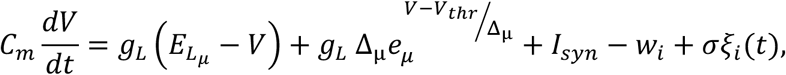

with *C*_*m*_ = 200 *pF* is the membrane capacity, *g*_*L*_ = 10 *nS* the leakage conductance, *E*_*Lμ*_ = −63 *mV*, −65 *mV* for *μ* = *E*, *I* the cell-type-dependent leakage reversal potential, Δ_μ_ = 2 *mV*, 0.5 *mV* and Δ_I_ = 0.5 *mV* the cell-type-dependent sodium sharpness, *V*_*thr*_ = −50 *mV* the spiking threshold, and *σ* = 0.0315 *pA* the amplitude of the noise with *ξ*_*i*_(*t*) random zero-mean Gaussian i.i.d. noise. *I*_*syn*_ the voltage-dependent input synaptic current of the neuron, that varies according to

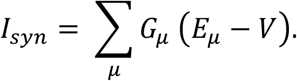

*G*_*μ*_ is the time-dependent synaptic conductance modelled by the exponential function

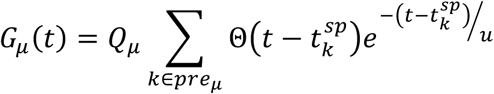

Which decays at a rate controlled by time constant *u* = 5 *ms*, and is incremented by a cell-type-dependent constant *Q*_*μ*_ = 1.5 *nS*, 5 *nS* across all brain regions for *μ* = *E*, *I* at each time 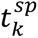 a spike is produced by neuron *k*. The sum is computed over all presynaptic neurons of type *μ*, and Θ(·) is the Heaviside function (1 when its argument is positive and 0 otherwise). The spike-frequency adaptation *w*_*i*_, that provides activity-dependent self-inhibition to the neuron, follows the differential equation

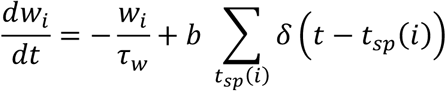

Summing over all spike times *t*_*sp*_(*i*) of neuron *i*, with *τ*_w_ = 500 *s* the adaptation decay time constant. Adaptation strength *b* = 0 *nS* throughout most of the simulations, unless stated otherwise, to model awake, high-neuromodulation states.

### Mean-field model

To bridge biophysical single-neuron dynamics to population firing rates at the brain region level without the prohibitive computational cost of simulating one equation for each of the many neurons within any region, we use a mean-field model of AdEx neural networks. Here, the mean-field model captures the firing rate of a mean neuron of any given type in a neural network as a response to the firing rates of other mean neurons of its own type and of all other types within the network. In the master equation formalism ^47^, this means that the firing rate *r*_*μ*_ of the mean neuron of type *μ* in an AdEx network is given by

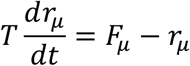

Where *x*_*μ*_ is the vector of firing rates for population *μ* ∈ {*E*, *I*} for each simulated network node and *T* is a time constant. The transfer function *F*_*μ*_(·), which relates each population’s output rate to the input it receives from itself and other populations is a cell-type-dependent polynomial whose parameters were fitted to reproduce the input-output relations of integrate-and-fire neurons in spiking simulations ^47^.

### Inter-regional coupling in TVB

To couple multiple regional mean field models together, we can write the previous differential equation as follows:

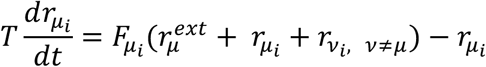

For a mean neuron of type *μ* in region *i*, with *v* is the other cell type than *μ* and 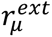 the external input from other brain regions i.e.

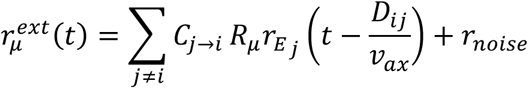

Where E neurons project synaptically onto both E and I neurons in other regions, with the ratio between *R*_*E*_ = 1 and varying *R*_*I*_ controlling relative projection strengths to E and to I neurons across regions. The directed, weighted graph of inter-regional synaptic connections *C*_j→*i*_ was taken from a 426-region tracer-derived structural connectome from the Allen Mouse Brain Institute ^33^. Inter-regional synaptic delays were modelled based on the matrix of tract lengths *D*_*i*j_ from the same data ^33^, with axonal propagation speed *v*_*ax*_ = 3*m*/*s*. Synaptic weights and delays were set as equal from both brain hemispheres.

### DMN emergence analysis

All simulations whose analyses are shown in figures from this point on are with a ratio between E-to-E and E-to-I inter-regional coupling strengths of *R* = 1.4.

### Insula, Cg, and PrL stimulation

Our primary analysis focuses on the insula to account for experimental findings reporting DMN suppression as observed with fMRI in response to optogenetic stimulation of the right anterior insula. We focus our stimulation on the agranular insular cortex rather than the granular insular cortex due to its stronger connectivity with limbic, autonomic, and higher-order cognitive regions ^48^. Unlike granular insular cortex, which primarily processes sensory inputs, AI is more integrative, playing a critical role in emotion, decision-making, and interoception. Evidence suggests that the mouse AI can play a major role in modulating neural dynamics across brain networks, as it serves as a structural outflow hub as opposed to the RSC node of the DMN which constitutes an inflow hub of the mouse structural connectome ^35^. Targeting AI allows us to better understand its contribution to brain network dynamics and neural processing and link to prior optogenetic stimulation studies.

For comparison, we consider the Cg and PrL as control stimulation sites. This allows us to compare salience circuit stimulation with within-DMN stimulation on the DMN and particularly its RSC node. As evidence for the RSC’s role as a structural ^35^ and functional ^24,26^ hub anchoring the rodent DMN has been provided by prior literature, it is crucial to understand causal influences of within-DMN as compared to salience circuit modulations on RSC dynamics.

To study the impact of stimulation on DMN dynamics, stimuli were square-wave pulses of firing rate of amplitude 0.1 Hz delivered for 0.5 s every 2 s for 150 s to match optogenetic protocols ^4^ after an initial transient of 5 s to all three subdivisions of the right agranular insula. The same procedure was repeated with the right Cg subdivisions together (Fig. 3) and separately (Fig. S2) and the right PrL stimulated instead of the insula. This procedure was repeated 100 times with different random seeds yield distinct initial conditions as well as random noise realizations.

The resulting simulated time series were averaged separately for each seed over stimulus presentations, i.e. over 2s long windows in which each stimulus was presented after 1s. For plotting, the trial-averaged evoked response was further downsampled to 10 Hz and averaged separately over all subdivisions of the RSC, all subdivisions of the Cg, and all subdivisions of the PrL.

### Stimulation to sensory and motor regions

To further compare insula stimulation effects to other regional stimulation effects, we successively stimulated every region in the right hemisphere, for a total of 171 stimulation sites in our model. For each stimulation site, the same procedure was repeated as for the insula, Cg, and PrL stimulation described above. This was repeated for 20 random realizations for each site. When a region contained multiple subdivisions, they were stimulated at the same time and amplitude.

### Local loss of I synaptic conductance

To study the impact of regional loss of I synaptic conductance, we modified the *Q*_*I*_ parameter in all subregions of the RSC bilaterally, then separately in all subregions of the Cg bilaterally, then separately in the PrL bilaterally. For each altered region, *Q*_*I*_ took the values 4 nS and 4.5 nS. In all other, unchanged regions, *Q*_*I*_ was maintained to the value of 5 nS. For each altered region and value of local *Q*_*I*_, 100 simulations were run with different random seeds. In each simulation, the insula was stimulated as described before. Analyses and statistics were performed in the same way as for previous sections.

### Hierarchical clustering analysis

To evaluate how robustly the DMN and frontal networks emerged as distinct networks in the absence of stimuli, we performed a clustering analysis. All clustering analyses were performed from Pearson correlation matrices averaged over 20 random realizations of 100s-long simulated time series. We used 1 – Pearson correlation as the distance metric and applied hierarchical clustering with complete linkage.

This was applied with brain-wide *Q*_*E*_ = 1.5 *nS*, *Q*_*I*_ = 5 *nS*. Keeping adaptation strength fixed at zero, we first varied the E-to-I/E-to-E ratio across values *R* = 1, 1.2, 1.4, 1.6. Then, we fixed *R* = 1.4 and varied adaptation strength across values *b* = 0, 25,50,100 *nS*.

### Parameter exploration

To characterize regions of parameter space associated with DMN suppression and enhancement in response to insula stimulation, we performed a parameter exploration varying synaptic conductance *Q*_*E*_ and *Q*_*I*_ in the RSC as well as the E-to-I/E-to-E ratio *R* and the spike-frequency adaptation strength *b* in the entire simulated brain. For each parameter combination, we performed 20 random realizations of 35-second simulated time series with insula stimulation every 2 seconds. We chose to vary *Q*_*E*_ and *Q*_*I*_ in the RSC due to the widespread effects observed in **Fig. 4** and the literature discussing the RSC as an anchor to the rodent DMN ^4,26^. In our parameter exploration, *Q*_*E*_ spanned values between 0.1 and 4.1 nS in steps of 0.2 nS while *Q*_*I*_spanned values between between 4 and 6 nS in steps of 0.1 nS.

We also varied the E-to-I/E-to-E ratio *R* as we found it to modulate DMN suppression (**Fig. S1**), by repeating simulations for *R* = 1, 1.2, 1.4, 1.6 successively. Spike-frequency adaptation was varied to simulate cholinergic modulation, which is highest at low adaptation in aroused states where the DMN can be suppressed by salient cues – adaptation strength took values *b* = 0, 25,50,100 *nS* in our simulations.

### Statistical testing

To evaluate the significance of suppression and enhancement in response to stimulation, we considered the difference between the trial-averaged, time-averaged evoked activity in the second before and in that after the stimulus in each trial. This difference was computed in each of the 100 independent random realizations of the model. Statistical testing compared this difference across independent realizations to zero with a two-tailed one-sample Wilcoxon test.

To compare two sets of stimulation responses, either for two different stimulation targets or for two different levels of inhibitory conductance, simulations were conducted with the same 100 random seeds across both sets of simulations. For each random seed, a pair of simulations with the same initial conditions and noise realization, differing only in the condition of interest i.e. stimulation site or inhibitory conductance, was obtained. This allowed us to compute differences between post- and pre-stimulus activity as before and perform a paired Wilcoxon signed-rank test. FDR correction for multiple comparisons was applied over the tests on signals from all RSC, Cg, and PrL subdivision considered.

### Code availability

All simulation codes will be made publicly available upon publication.

## Acknowledgements

This research was supported by the Extramural Research Programs of U.S. NIH: NIMH (R01MH126518) and NINDS (RF1NS086085).

## Author contributions

T.-A. E. N. and V. M. designed research, T.-A. E. N. performed research, T.-A. E. N. and V. M. analyzed the data and wrote the paper.

## Supplementary Materials

**Figure S1.**
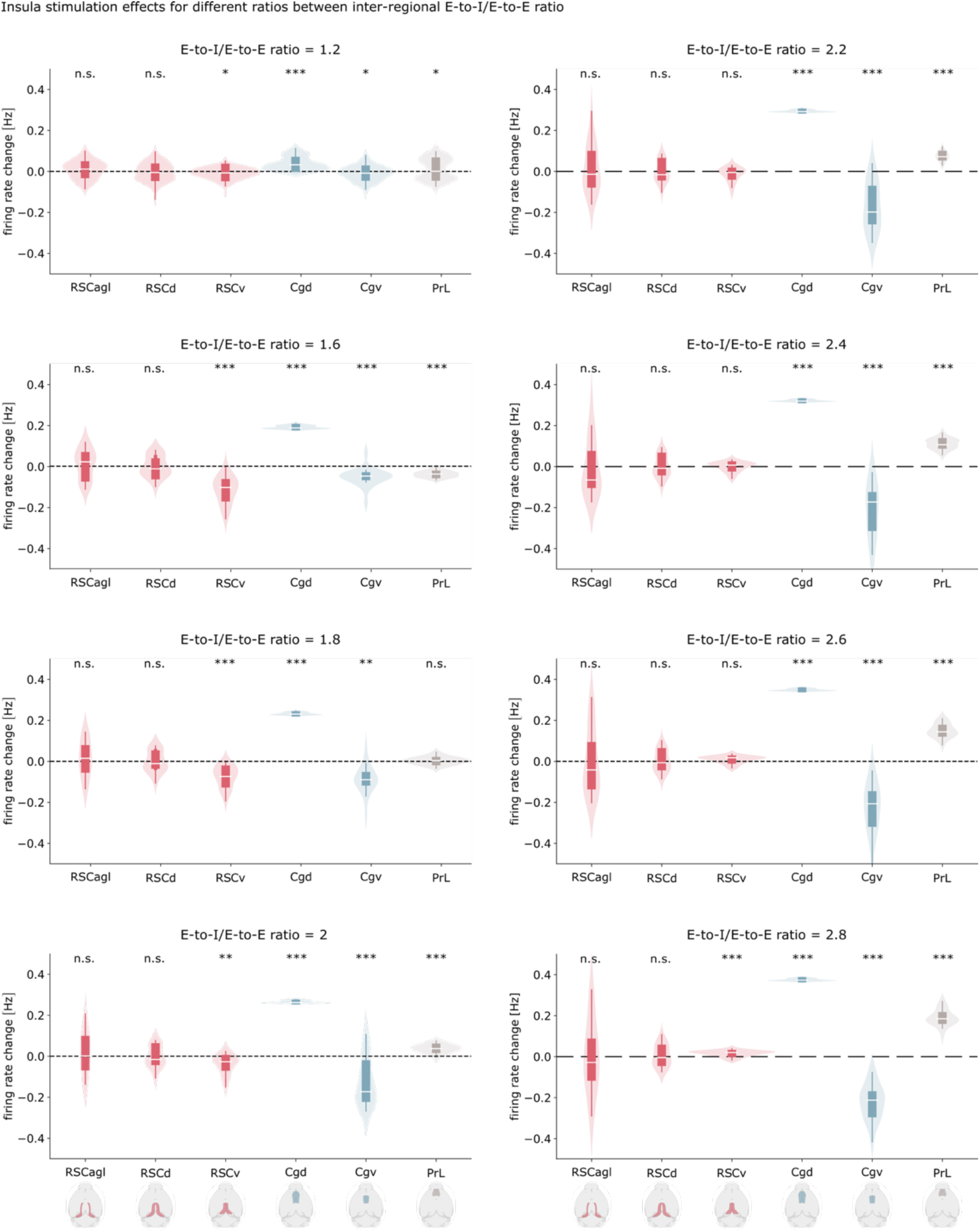
Insula stimulation produces coordinated DMN suppression across a range of ratios between E-to-E and E-to-I inter-regional projection weights. The RSC is found to be suppressed RSCv across ratios from 1.2 to 2.2, the Cgv across ratios from 1.2 to 2.8, and the PrL across ratios from 1.2 to 1.8. The results show that DMN suppression is robustly obtained ratios provided stronger E-to-I than E-to-E weights.

**Figure S2.**
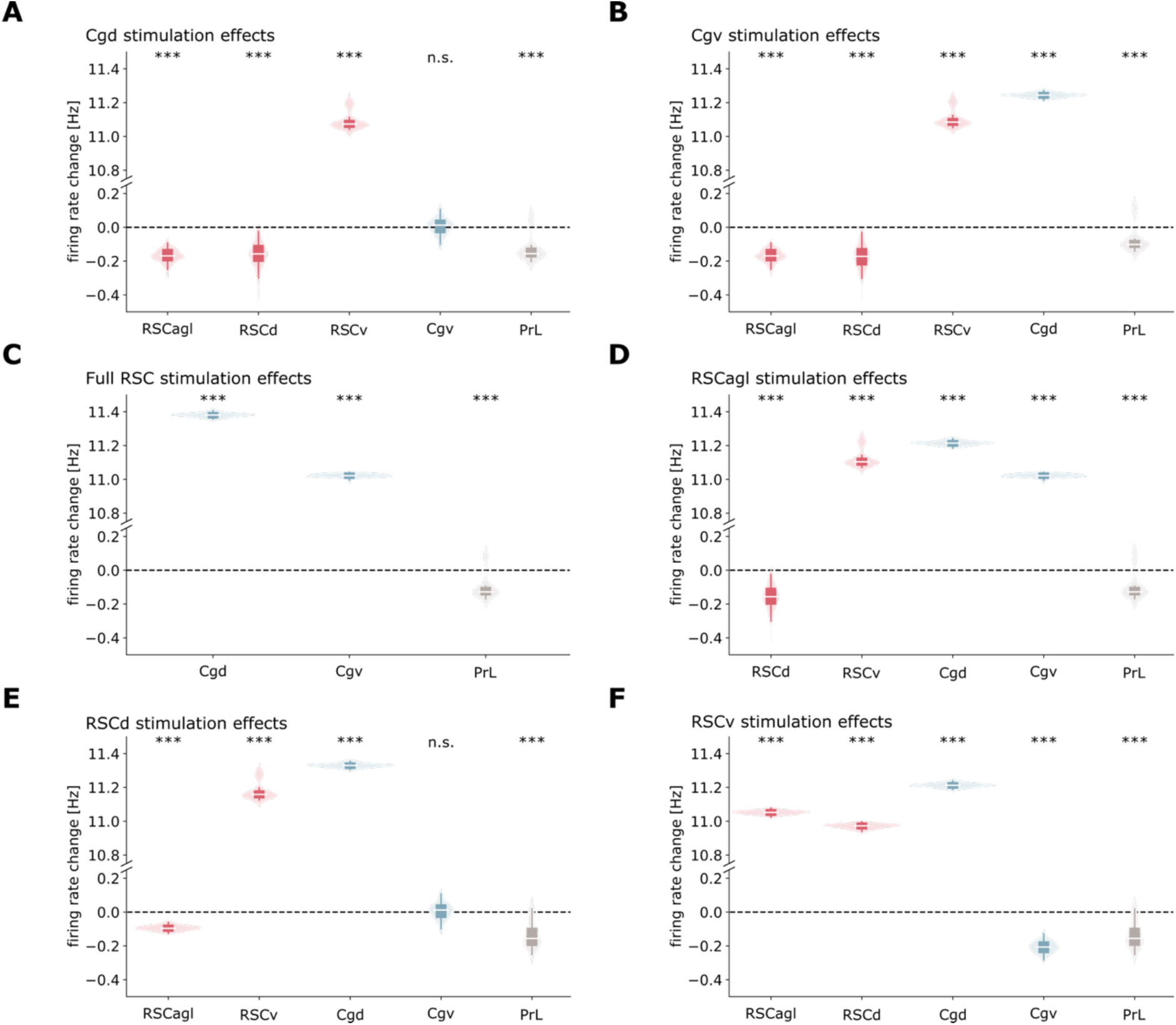
Effects of RSC and Cg subdivision stimulation on DMN activity. **(A-B)** Stimulation of the Cgd **(A)** and Cgd **(B)** both enhance RSCv dynamics while suppressing other RSC subdivisions as well as the PrL. **(C-F)** Stimulation of the full RSC **(C)** as well as of its RSCagl **(D)**, RSCd **(E)**, and RSCv **(F)** subdivisions separately all enhance the Cgd and suppress the PrL. Effects on the Cgv are subdivision-specific, with net enhancement from full RSC stimulation. The results show the Cg and RSC, when stimulated, mutually enhance each other, unlike the insula when stimulated (Figure 2).

**Figure S3.**
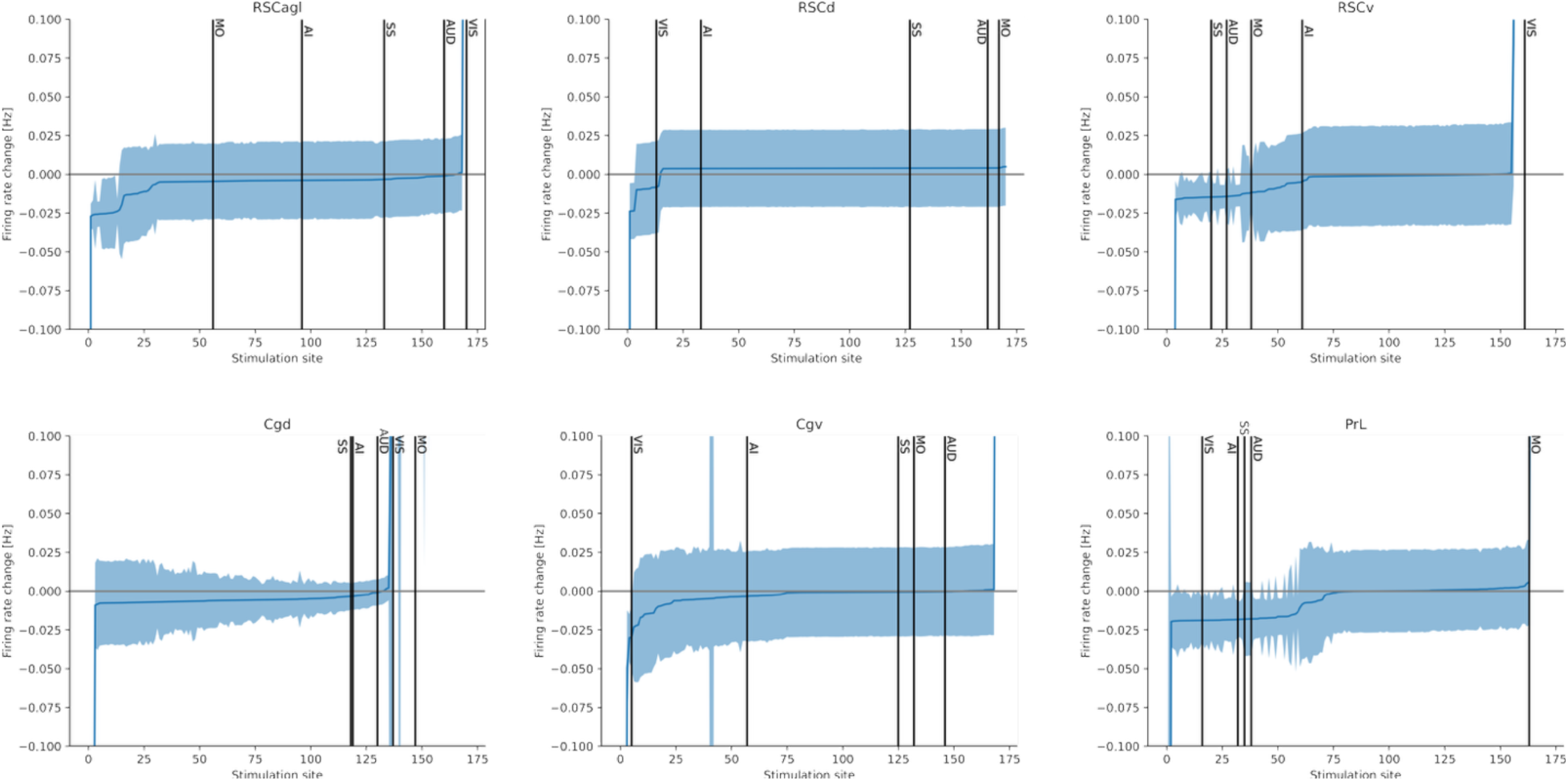
Effects of stimulation to each brain region on DMN activity. The simulated firing rate change in each brain region is shown as a function of the stimulation site. Different panels show firing rate change in different DMN subdivisions. Stimulation sites spanned all regions of the right hemisphere and are ranked in increasing order (from most negative to most positive) of firing rate change in each DMN subdivision and each panel. Black vertical lines indicate stimulation to the auditor (AUD), somatosensory (SS), visual (VIS), and motor (MO) cortex.

**Figure S4.**
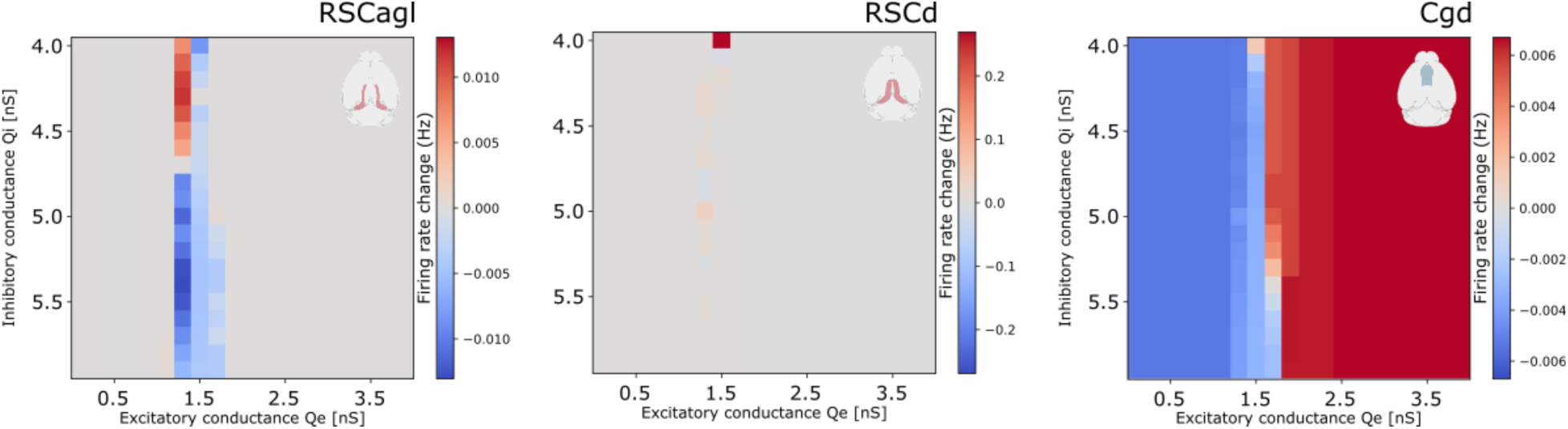
Parameter exploration of local E-I balance impacts on agranular and dorsal DMN subdivision suppression. Heatmaps of simulated subregional response to insula stimulation in the RSCagl (left), RSCd (middle), and Cg (right) as a function of E and I synaptic conductance in the RSC. The results show the region-specific and distributed effects of E-I imbalance in the RSC.

**Figure S5.**
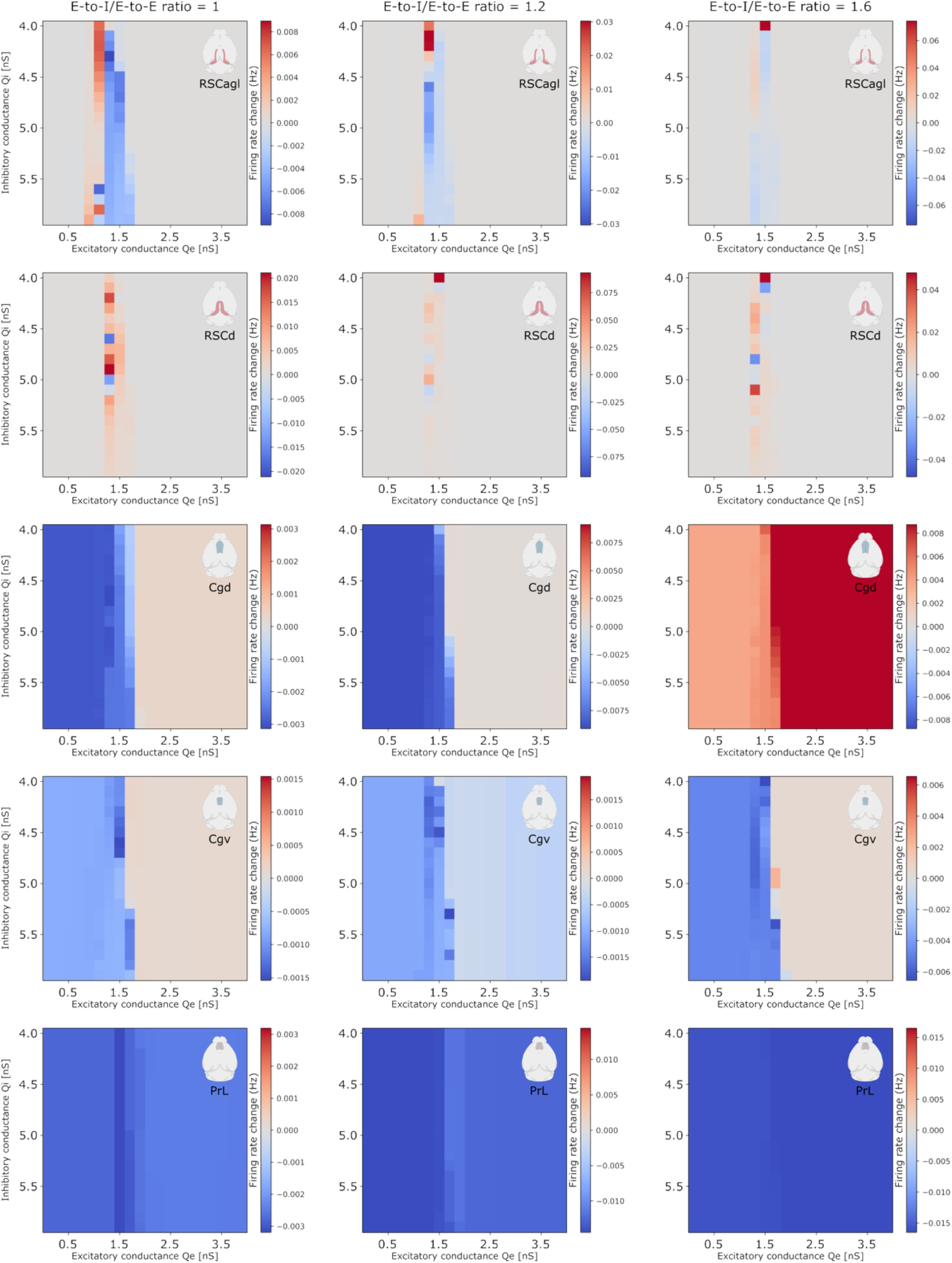
Effect of E-to-I/E-to-E ratio on DMN-wide suppression. Heatmaps of simulated responses in all DMN subregions except the RSCv, as a function of E and I synaptic conductance *Q*_*E*_ and *Q*_*I*_ in the RSC for a varying E-to-I/E-to-E ratio and zero spike-frequency adaptation.

**Figure S6.**
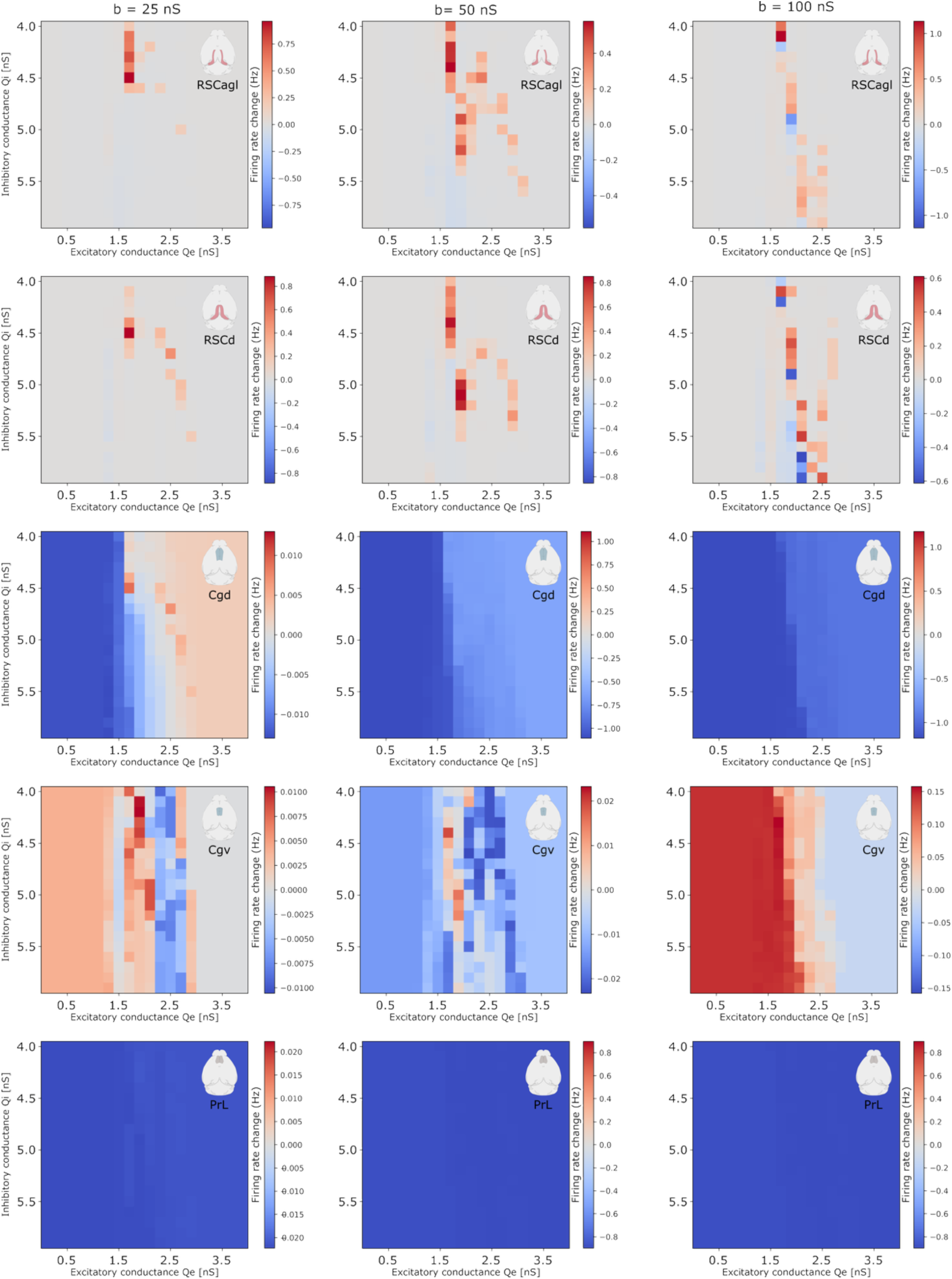
Effect of cholinergic neuromodulation via spike-frequency adaptation on DMN-wide suppression. Heatmaps of simulated responses in all DMN subregions except the RSCv, as a function of E and I synaptic conductances *Q*_*E*_ and *Q*_*I*_ in the RSC for varying levels of modeled cholinergic modulation via spike-frequency adaptation *b* and a fixed E-to-I/E-to-E ratio of 1.4.

**Figure S7.**
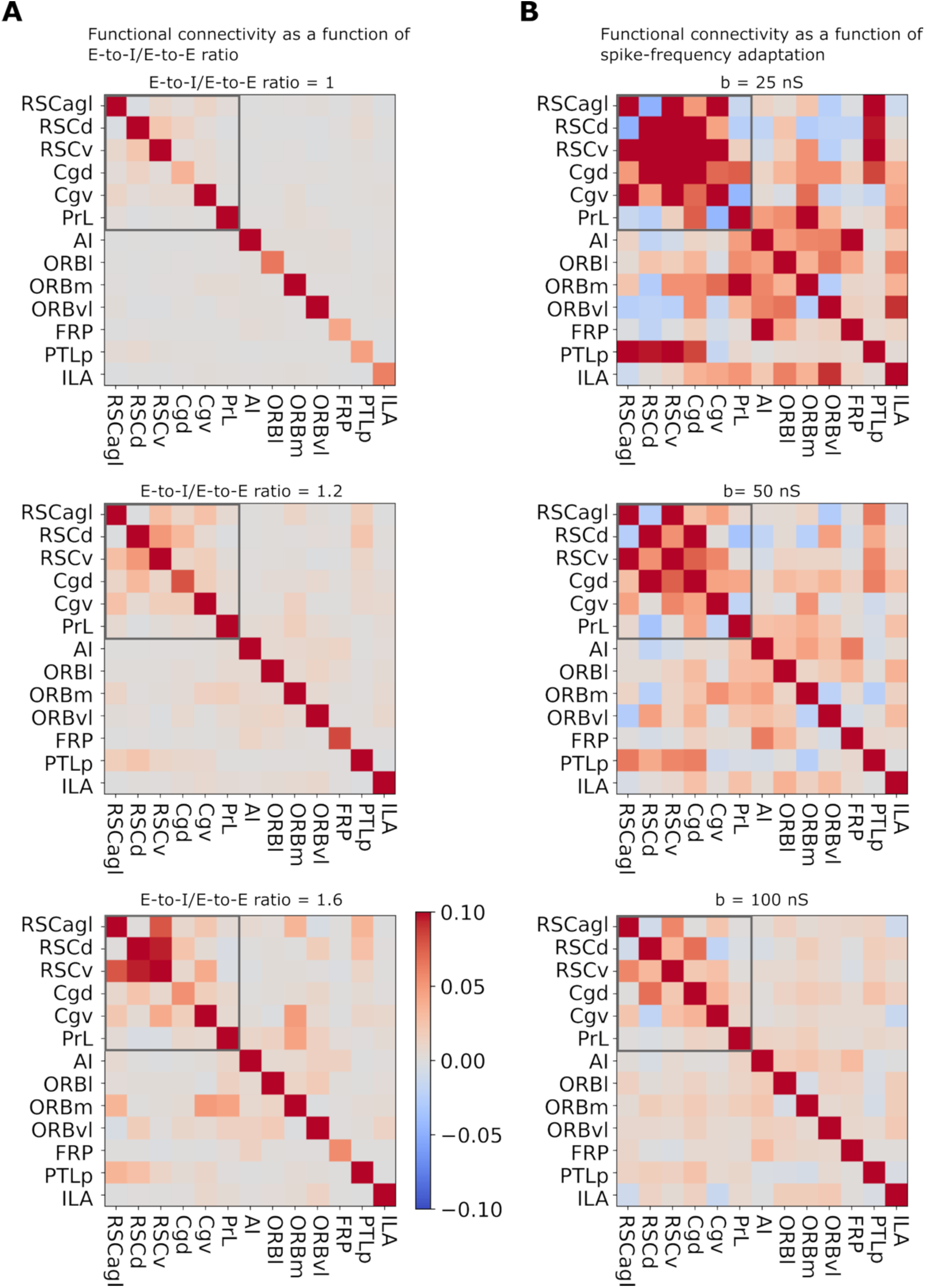
Resting-state functional connectivity matrices between the DMN, insula, and putative FN for different values of E-to-I/E-to-E ratio (**A**) and spike-frequency adaptation (**B**) model parameters. The results show intra-DMN functional connectivity growing with E-to-I/E- to-E ratio, as well as intra-DMN negative functional connectivity emerging in the presence of spike-frequency adaptation.

**Figure S8.**
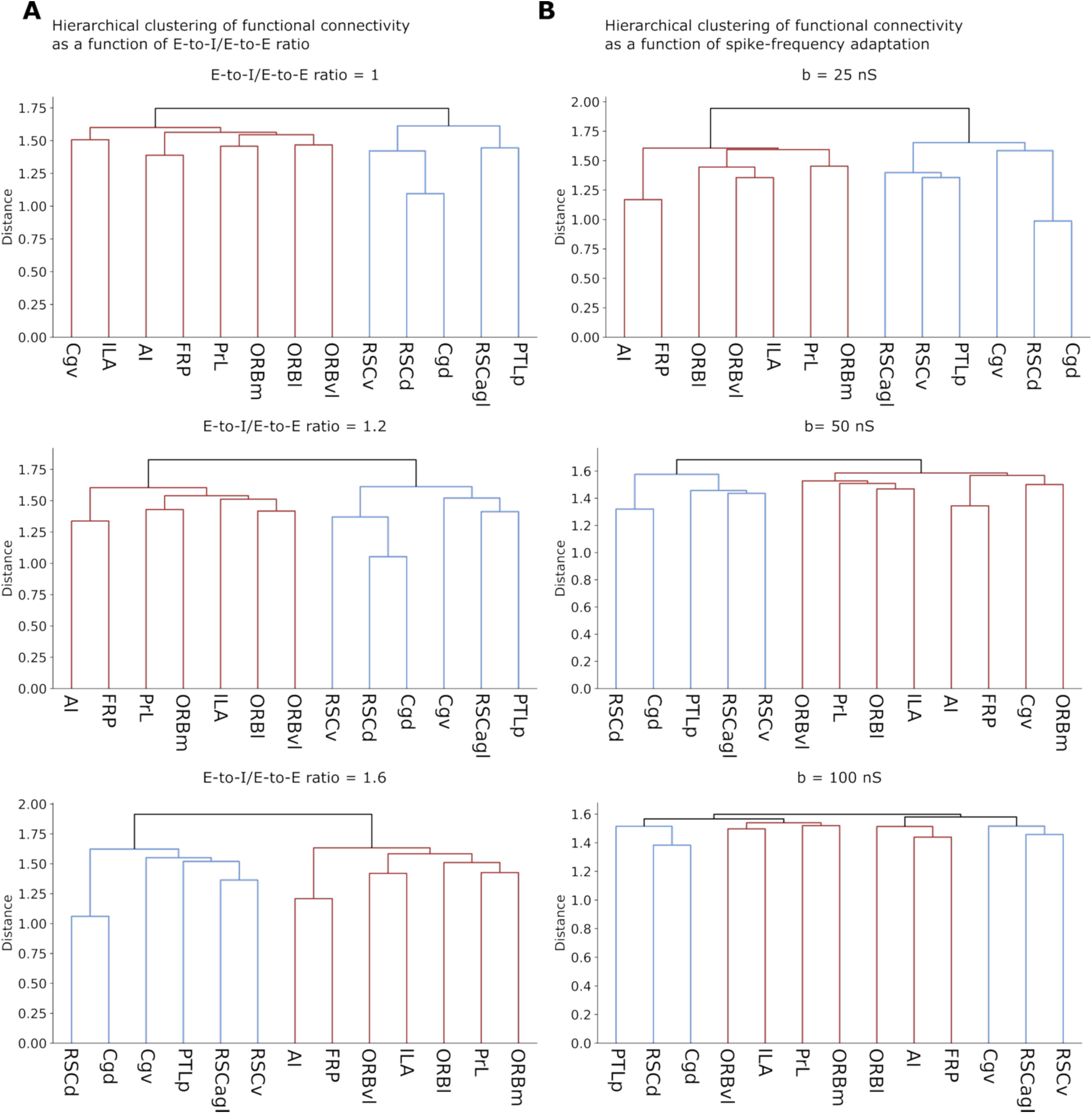
Resting-state functional connectivity hierarchical clustering dendrograms for different values of E-to-I/E-to-E ratio (**A**) and spike-frequency adaptation (**B**) model parameters. The results show the robustness of DMN emergence (in blue) across simulation parameters, as well as DMN breakdown at high spike-frequency adaptation.

**Table S1.** Three distinct DMN breakdown modes. Our analysis revealed that simulated DMN suppression fails through three mechanistically distinct modes: loss of responsiveness, reversal from suppression to enhancement, and network fragmentation.

| Feature | Parameter Range (local) | Parameter Range Global (EI to EE) | Parameter Range Adaptation | DMN Behavior | Interpretation |
| --- | --- | --- | --- | --- | --- |
| **DMN suppression** | $Q_E = 1.3$ to $1.5$ nS,<br>$Q_I > 5$ nS,<br>Or $Q_E = 1.5$,<br>$Q_I > 4$ nS | Ratio $> 1$ | $b < 100$ nS | Suppression in RSCv, Cgv, and PrL | Specific E and I conductance ranges are needed to simulate DMN suppression |
| **Loss of RSC response, DMN response fragmentation** | $Q_E = 1.4$ to $1.5$ nS,<br>$Q_I < 5$ nS | Ratio $> 1$ | $b < 100$ nS | Suppression in Cgv and PrL, no response in RSCv | Specific E and I conductance ranges are needed to simulate DMN suppression |
| | $Q_E < 0.7$ or $Q_E > 1.7$ nS | Ratio $\geq 1$ | $b = 0$ nS | | Specific E and I conductance ranges are needed to simulate DMN suppression |
| | $Q_E = 1.3$ to $1.5$ nS,<br>$Q_I > 5$ nS,<br>Or $Q_E = 1.5$,<br>$Q_I > 4$ nS | Ratio $= 1$ | $b = 0$ nS | | DMN suppression is robust to ratio provided ratio $> 1$ |
| | $Q_E = 1.3$ to $1.5$ nS,<br>$Q_I > 5$ nS,<br>Or $Q_E = 1.5$,<br>$Q_I > 4$ nS | Ratio $= 1.4$ | $b = 100$ nS | Suppression in PrL, no response in RSCv, enhancement in Cgv | High adaptation disrupts intra-DMN and DMN-insula interactions |
| **RSC reversal, DMN response fragmentation** | $Q_E = 0.7$ to $1.4$ nS, $Q_I > 4$ nS | Ratio $= 1.4$ | $b = 0$ nS | Suppression in PrL and Cg, enhancement in RSCv | Low activity due to low excitability, promoting enhancement over suppression by stimulation |
| | $Q_E = 1.6$ nS, $Q_I = 4.9$ to $5.5$ nS | Ratio $= 1.4$ | $b = 0$ nS | Suppression in PrL, enhancement in RSCv and Cgv | High excitability leads to distributed enhancement |
| | $Q_E > 1.5$ nS, $Q_I > 4$ nS | Ratio $= 1$ | $b > 0$ nS | Suppression in PrL, enhancement in RSCv, no consistent response in Cgv | Adaptation reduces firing rate, promoting enhancement over suppression by stimulation |

**Table S2.** Robustness of DMN emergence and parameter regimes for DMN integrity. DMN emergence as a distinct network is only disrupted at high spike-frequency adaptation rates. Although E-to-I/E-to-I inter-regional connectivity ratio influences functional connectivity in the DMN, it does not disrupt DMN integrity.

| Feature | Parameter Range Global (EI to EE) | Parameter Range Adaptation | DMN Behavior | Interpretation |
| --- | --- | --- | --- | --- |
| **DMN emergence as a network** | Ratio > 1 | $b < 100 \text{ nS}$ | DMN clusters as a network apart from insula and frontal network | DMN robustly emerges as a network across parameter regimes |
| **DMN reduced functional connectivity** | Ratio = 1 | $b = 0 \text{ nS}$ | DMN remains a separate cluster but with reduced internal functional connectivity | Sufficiently strong E-to-I connectivity maintaining global E-I balance is needed for strong intra-DMN connectivity |
| **DMN spontaneous fragmentation** | Ratio = 1.4 | $b = 100 \text{ nS}$ | DMN splits into two clusters: RSCv, RSCagl, and Cgv in one cluster, RSCd, Cgd, and PTLp in the other | Adaptation generates neural self-inhibition that gives rise to anticorrelation between DMN nodes |

## References

1 Menon, V. 20 years of the default mode network: A review and synthesis. Neuron 111, 2469–2487 (2023).

2 Seeley, W. W. et al. Dissociable intrinsic connectivity networks for salience processing and executive control. Journal of neuroscience 27, 2349–2356 (2007).

3 Sridharan, D., Levitin, D. J. & Menon, V. A critical role for the right fronto-insular cortex in switching between central-executive and default-mode networks. Proceedings of the National Academy of Sciences 105, 12569–12574 (2008).

4 Menon, V. et al. Optogenetic stimulation of anterior insular cortex neurons in male rats reveals causal mechanisms underlying suppression of the default mode network by the salience network. Nature Communications 14, 866 (2023).

5 Greicius, M. D., Krasnow, B., Reiss, A. L. & Menon, V. Functional connectivity in the resting brain: a network analysis of the default mode hypothesis. Proceedings of the National Academy of Sciences 100, 253–258 (2003).

6 Molnar-Szakacs, I. & Uddin, L. Q. Self-processing and the default mode network: interactions with the mirror neuron system. Frontiers in human neuroscience 7, 571 (2013).

7 Menon, V. Large-scale brain networks and psychopathology: a unifying triple network model. Trends in cognitive sciences 15, 483–506 (2011).

8 Goodkind, M. et al. Identification of a common neurobiological substrate for mental illness. JAMA psychiatry 72, 305–315 (2015).

9 McTeague, L. M. et al. Identification of common neural circuit disruptions in cognitive control across psychiatric disorders. American Journal of Psychiatry 174, 676–685 (2017).

10 Padmanabhan, A., Lynch, C. J., Schaer, M. & Menon, V. The default mode network in autism. Biological Psychiatry: Cognitive Neuroscience and Neuroimaging 2, 476–486 (2017).

11 Weissman, D. H., Roberts, K., Visscher, K. & Woldorff, M. The neural bases of momentary lapses in attention. Nature neuroscience 9, 971–978 (2006).

12 Menon, V., Anagnoson, R., Mathalon, D., Glover, G. & Pfefferbaum, A. Functional neuroanatomy of auditory working memory in schizophrenia: relation to positive and negative symptoms. Neuroimage 13, 433–446 (2001).

13 Kwon, H., Reiss, A. L. & Menon, V. Neural basis of protracted developmental changes in visuo-spatial working memory. Proceedings of the National Academy of Sciences 99, 13336–13341 (2002).

14 Garrity, A. G. et al. Aberrant “default mode” functional connectivity in schizophrenia. American journal of psychiatry 164, 450–457 (2007).

15 Zhou, H.-X. et al. Rumination and the default mode network: Meta-analysis of brain imaging studies and implications for depression. Neuroimage 206, 116287 (2020).

16 Rubenstein, J. L. R. & Merzenich, M. M. Model of autism: increased ratio of excitation/inhibition in key neural systems. Genes Brain and Behavior 2, 255–267 (2003). 10.1034/j.1601-183X.2003.00037.x

17 Nelson, S. B. & Valakh, V. Excitatory/inhibitory balance and circuit homeostasis in autism spectrum disorders. Neuron 87, 684–698 (2015).

18 Supekar, K., Ryali, S., Mistry, P. & Menon, V. Aberrant dynamics of cognitive control and motor circuits predict distinct restricted and repetitive behaviors in children with autism. Nature Communications 12, 3537 (2021).

19 Liu, Y. et al. A selective review of the excitatory-inhibitory imbalance in schizophrenia: underlying biology, genetics, microcircuits, and symptoms. Frontiers in cell and developmental biology 9, 664535 (2021).

20 Canitano, R. & Pallagrosi, M. Autism spectrum disorders and schizophrenia spectrum disorders: excitation/inhibition imbalance and developmental trajectories. Frontiers in psychiatry 8, 69 (2017).

21 Antoine, M. W., Langberg, T., Schnepel, P. & Feldman, D. E. Increased excitation-inhibition ratio stabilizes synapse and circuit excitability in four autism mouse models. neuron 101, 648–661. e644 (2019).

22 Yizhar, O. et al. Neocortical excitation/inhibition balance in information processing and social dysfunction. Nature 477, 171–178 (2011).

23 Antoine, M. W., Langberg, T., Schnepel, P. & Feldman, D. E. Increased Excitation-Inhibition Ratio Stabilizes Synapse and Circuit Excitability in Four Autism Mouse Models. Neuron 101, 648–+ (2019). 10.1016/j.neuron.2018.12.026

24 Gutierrez-Barragan, D. et al. Unique spatiotemporal fMRI dynamics in the awake mouse brain. Current biology 32, 631–644. e636 (2022).

25 Chao, T.-H. H. et al. Neuronal dynamics of the default mode network and anterior insular cortex: Intrinsic properties and modulation by salient stimuli. Science Advances 9, eade5732 (2023).

26 Nghiem, T.-A. E. et al. Space wandering in the rodent default mode network. Proceedings of the National Academy of Sciences 121, e2315167121 (2024).

27 Sanz Leon, P., et al. The Virtual Brain: a simulator of primate brain network dynamics. Frontiers in neuroinformatics 7, 10 (2013).

28 Sanz-Leon, P., Knock, S. A., Spiegler, A. & Jirsa, V. K. Mathematical framework for large-scale brain network modeling in The Virtual Brain. Neuroimage 111, 385–430 (2015).

29 Schirner, M., McIntosh, A. R., Jirsa, V., Deco, G. & Ritter, P. Inferring multi-scale neural mechanisms with brain network modelling. elife 7, e28927 (2018).

30 Zimmermann, J. et al. Differentiation of Alzheimer’s disease based on local and global parameters in personalized Virtual Brain models. NeuroImage: Clinical 19, 240–251 (2018).

31 Kringelbach, M. L., Perl, Y. S., Tagliazucchi, E. & Deco, G. Toward naturalistic neuroscience: Mechanisms underlying the flattening of brain hierarchy in movie-watching compared to rest and task. Science Advances 9, eade6049 (2023).

32 Goldman, J. S., et al. A comprehensive neural simulation of slow-wave sleep and highly responsive wakefulness dynamics. bioRxiv (2021).

33 Oh, S. W. et al. A mesoscale connectome of the mouse brain. Nature 508, 207–214 (2014).

34 Maier-Hein, K. H. et al. The challenge of mapping the human connectome based on diffusion tractography. Nature communications 8, 1349 (2017).

35 Coletta, L. et al. Network structure of the mouse brain connectome with voxel resolution. Science advances 6, eabb7187 (2020).

36 Fox, M. D. et al. The human brain is intrinsically organized into dynamic, anticorrelated functional networks. Proceedings of the National Academy of Sciences 102, 9673–9678 (2005).

37 Vogt, B. A. Cingulate cortex in the three limbic subsystems. Handbook of clinical neurology 166, 39–51 (2019).

38 Menon, V. & Uddin, L. Q. Saliency, switching, attention and control: a network model of insula function. Brain structure and function 214, 655–667 (2010).

39 Lynch, C. J. et al. Default mode network in childhood autism: posteromedial cortex heterogeneity and relationship with social deficits. Biological psychiatry 74, 212–219 (2013).

40 Gozzi, A. & Schwarz, A. J. Large-scale functional connectivity networks in the rodent brain. Neuroimage 127, 496–509 (2016).

41 Tsai, P.-J. et al. Converging structural and functional evidence for a rat salience network. Biological psychiatry 88, 867–878 (2020).

42 Lyamzin, D. & Benucci, A. The mouse posterior parietal cortex: Anatomy and functions. Neuroscience research 140, 14–22 (2019).

43 Pagani, M. et al. Biological subtyping of autism via cross-species fMRI. bioRxiv (2025).

44 Pagani, M. et al. mTOR-related synaptic pathology causes autism spectrum disorder-associated functional hyperconnectivity. Nature communications 12, 6084 (2021).

45 Zerbi, V. et al. Brain mapping across 16 autism mouse models reveals a spectrum of functional connectivity subtypes. Molecular psychiatry 26, 7610–7620 (2021).

46 Ding, X., Froudist-Walsh, S., Jaramillo, J., Jiang, J. & Wang, X.-J. Cell type-specific connectome predicts distributed working memory activity in the mouse brain. elife 13, e85442 (2024).

47 Di Volo, M., Romagnoni, A., Capone, C. & Destexhe, A. Biologically realistic mean-field models of conductance-based networks of spiking neurons with adaptation. Neural computation 31, 653–680 (2019).

48 Menon, V. Insular cortex: A hub for saliency, cognitive control, and interoceptive awareness. (2025).

